# ‘Digital Reprogramming’ Decodes Epigenetic Barriers of Cell Fate Changes

**DOI:** 10.1101/2025.01.22.634227

**Authors:** Ana Janeva, Christopher A. Penfold, Sara Llorente-Armijo, Huiwen Li, Tomas Zikmund, Marco Stock, Jerome Jullien, Tobias Straub, Ignasi Forne, Axel Imhof, Juan M. Vaquerizas, John B. Gurdon, Eva Hörmanseder

**Author notes:** corresponding authors; correspondence to E.H., A.J. contributed equally (Ana Janeva, Christopher A. Penfold).

## Abstract

The fates of differentiated cells in our body can be induced to change by nuclear reprogramming. In this way, cells valuable for therapeutic purposes and disease modeling can be produced. However, the efficiency of this process is low, partly due to properties of somatic donor nuclei which stabilize their differentiated fate but also act as barriers reprogramming-associated cell fate changes. The identity of these reprogramming barriers is not fully understood.

Here, we developed an artificial intelligence-based approach to model nuclear reprogramming and used it to identify the chromatin modification H3K27ac as an epigenetic barrier to reprogramming-induced cell fate changes. Using reprogramming by nuclear transfer (NT) to eggs of *Xenopus laevis* as a model system, we profiled chromatin modifications in differentiated cell types alongside gene expression patterns before and after reprogramming. Our model integrated the data and by leveraging model predictions, we find that genes resisting inactivation during reprogramming display chromatin modification barcodes. This revealed H3K27ac as a novel candidate barrier to NT reprogramming. Reducing H3K27ac levels using p300/CBP inhibitors before reprogramming led to an improved downregulation of genes linked to H3K27ac-modified enhancers after reprogramming. Importantly, these effects were accompanied by improved embryonic development of the resulting nuclear transfer embryos.

In summary, our study identified H3K27ac as a safeguarding mechanism of cellular identities and as a reprogramming barrier during NT. Hence, the here-developed ‘Digital Reprogramming” approach is capable of modelling and improving current cell-fate reprogramming strategies.

## INTRODUCTION

The fusion of gametes results in a totipotent zygote capable of producing all cell types of an organism. As the zygote undergoes cell divisions and gradually restricts its potency, cell identities emerge and are stably passed on to the daughter cells. Developmental cues are integrated, and specific transcription factor networks are acquired alongside a complex set of epigenetic traits that stabilize their cellular identity with each step of lineage commitment. Thus, once a cell fate is established in a healthy organism, it rarely changes, and such abnormal changes are often associated with cell death or disease. However, several experimental manipulations, such as transcription factor (TF) overexpression^1^, cell-cell fusion^2^, or somatic cell nuclear transfer (SCNT)^3^ can revert cells to an undifferentiated state and alter their identity.

The process of SCNT, which involves the transplantation of a somatic nucleus into an enucleated vertebrate egg, can reprogram differentiated nuclei to totipotency and give rise to all cell types of a cloned organism^3^. However, this process is inefficient, as cloned embryos have a poor survival rate^4^ and face many developmental abnormalities. It has been hypothesized that a transcriptional memory of the somatic cell fate, which stabilizes cellular identities during normal development, also prevents cell fate changes during reprogramming.

Indeed, previous studies have provided examples where the transcription state characteristic of the donor cell fate was inherited by most cell types of NT-embryos^5^ acting as an important barrier to efficient cell fate conversion^5–7^. Specifically, ectoderm cells in NT-embryos derived from endoderm donor cells continue to aberrantly express endoderm genes^8^. The expression of these genes represents a failure of the egg to downregulate the endoderm program correctly, conceived as a persistent transcriptional memory of the endoderm state (ON-memory). A second class of memory genes collectively represents those expressed in the target tissue type in *in vitro* fertilized (IVF) embryos, but not in the donor cell type, which fails to be activated in the NT-embryo, or is activated at lower levels than in IVF-embryos^8^. These genes are said to have inherited memory of the suppressed state of the donor cell type and are termed OFF-memory genes.

Mechanistic studies showed that a failure to activate the expression of OFF-memory genes may be caused by the persistence of repressive chromatin marks inherited from the donor cell type, suggestive of epigenetic memory of repressed chromatin states. In mice, aberrant OFF-memory effects at the two-cell stage have been linked to the persistence of the repressive mark H3K9me3^9^, which could be alleviated by *Kdm4a/d* overexpression^9–11^. Several other chromatin modifications have also been implicated in resistance to reprogramming (reviewed in^12^) including histone deacetylation^13–16^, DNA methylation^17–19^, and various mammal-specific features including extraembryonic defects^22^, and H3K27me3 imprinting defects^23–25^. High-throughput chimeric systems that use *Xenopus* germinal vesicle (GV) to reprogram mouse nuclei, suggest that genes may resist reprogramming by multiple epigenetic barriers and mechanisms^26^. The mechanisms underlying ON-memory in reprogrammed cells are less understood. In part, the persistence of trimethylation at histone H3 lysine K4 (H3K4me3) around the transcription start site (TSS) has been reported to maintain ON-memory in SCNT reprogramming^8^. Injection of the H3K4-specific demethylase Kdm5b can mitigate this barrier in *Xenopus*, resulting in increased reprogramming efficiency^8^, with similar results seen in mammalian systems^27–29^. These results indicate that active transcriptional states are memorized from the donor nucleus and could be passed on to the reprogrammed cell through epigenetic mechanisms of active chromatin states^5,6^.

However, in part due to the complex molecular nature of cell fate reprogramming, the molecular mechanisms underlying ON-memory, alongside OFF-memory are currently not fully understood. Specifically, it is unclear which ‘active’ chromatin marks alongside H3K4me3 contribute to ON-memory. Furthermore, it is unknown if active marks act in concert or in parallel with repressive marks, forming an “epigenetic barcode” that stabilizes cell fate-specific gene expression and prevents cell fate reprogramming. Novel computational approaches are missing which integrate complex multi-omics datasets generated during reprogramming to inform on the underlying biological mechanisms of reprogramming.

In the present study, we uncovered epigenetic features of differentiated cells that stabilize their cell identity and prevent nuclear reprogramming, with a focus on the memory of active chromatin states. We constructed a large-scale reference dataset of histone modifications and gene expression profiles in different tissue types, reprogrammed nuclei from two of those tissues, and catalogued memory class genes in the resulting NT-embryos. Leveraging these datasets, we developed an artificial intelligence approach based on convolutional neural networks (CNNs) and built a predictive model, termed Digital Reprogramming. Based on histone modification levels and other genomic features of the somatic donor cell types, Digital reprogramming was able to accurately predict transcriptome changes and memory status of genes during reprogramming. Using this approach, we further identified candidate reprogramming barriers linked to the inheritance of active gene expression states, or ON-memory. Digital Reprogramming highlights previously reported reprogramming barriers such as H3K4me3, demonstrating its accuracy and robustness. Importantly, Digital Reprogramming also identifies acetylation on histone H3 lysine 27 (H3K27ac) as a novel feature for a specific group of ON-memory genes. Interfering with the histone acetyltransferase p300/CBP to reduce H3K27ac levels results in a decreased number of genes with ON-memory status during reprogramming and, importantly, improved development of NT-embryos.

In summary, our Digital Reprogramming successfully predicts reprogramming outcomes and identifies roadblocks. Our results implicate H3K27ac as a protective mechanism of differentiated cell identities and a barrier to cell fate changes during reprogramming.

## RESULTS

### Construction of large-scale, multi-omic reference datasets of nuclear reprogramming

To build a computational model capable of predicting reprogramming outcomes and barriers, we first needed to generate reference datasets for training and testing such a model. Therefore, we first defined memory genes in different cell types of *Xenopus laevis* NT-embryos, and profiled histone modifications in two different donor cell types.

For this, donor mesoderm nuclei were transplanted to enucleated eggs to generate NT-embryos (**Figure 1a**). IVF-embryo were generated as controls for wild-type gene expression. At gastrula stage 11, endoderm tissues were isolated and subjected to RNA-seq. Then, differential gene expression analyses were performed between the donor-mesoderm, the NT-endoderm, and the IVF-endoderm tissue samples, resulting in 9890 differentially expressed genes (DEGs). To identify 1737 ON-memory genes, genes were selected that were expressed in the donor mesoderm tissue and that remained significantly upregulated in NT-endoderm, when compared to IVF-endoderm (**Figure 1 b,c,d**; see Methods section for cut-offs and filtering criteria). Instead, genes expressed in donor cells that were downregulated to comparable levels between NT and IVF revealed 3072 reprogrammed-down genes. Vice versa, 2013 mesoderm OFF-memory genes were identified (**Figure 1 b,c,d**) by intersecting genes expressed at lower levels in donor-mesoderm than in IVF-endoderm with genes that are still expressed at lower levels in NT-endoderm when compared to IVF-endoderm. Instead, intersection with genes expressed at similarly high levels in NT-endoderm as in IVF-endoderm revealed 3068 reprogrammed-up genes. Corresponding published endoderm-to-ectoderm reprogramming datasets^8^ were analyzed (**Figure S1a**) in the same way. This identified 1382 endoderm ON-memory genes and 1372 endoderm OFF-memory genes in endoderm-derived NT-ectoderm cells (**Figure S1 b,c**). Mesoderm ON-memory genes, in contrast to correctly reprogrammed-down genes, maintained aberrantly high expression levels in the endoderm of NT-embryos when compared to the endoderm of IVF-embryos. **(Figure 1c).** Similarly, endoderm ON-memory genes, in contrast to reprogrammed-down genes, maintained high expression levels in the ectoderm of NT-embryos in contrast to the ectoderm of IVF-embryos (**Figure S1c)**. We further observe that the number of identified genes with ON- and OFF-memory status differs depending on the donor and target tissue used (**Figure 1b and Figure S1b)**. Together, these results show that ON- and OFF-memory gene expression is a defining feature of nuclear reprogramming using NT.

**Figure 1:**
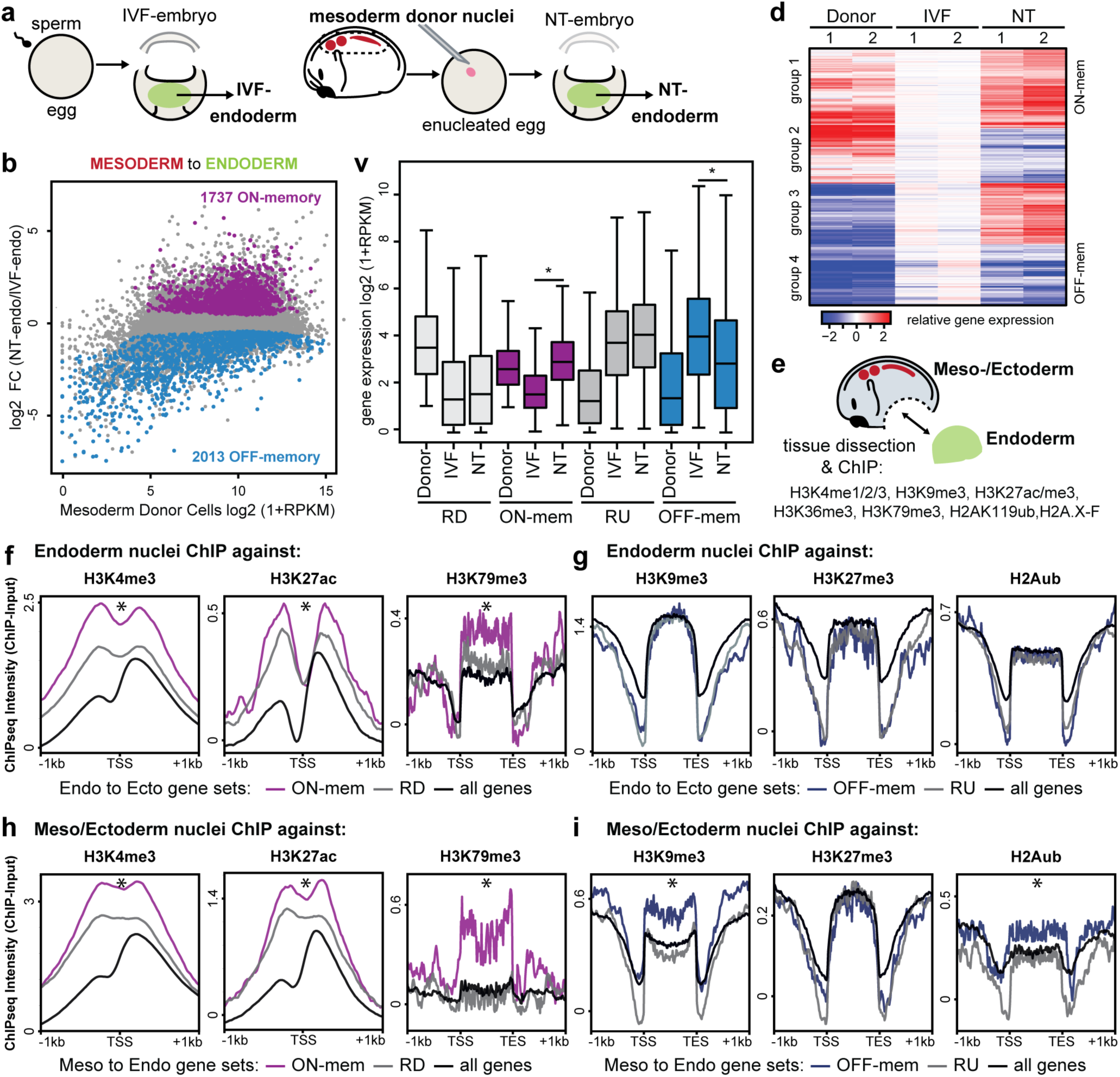
Memory genes resist reprogramming. **(a)** Memory genes were identified in stage 11 endoderm following somatic cell nuclear transfer (SCNT) with mesoderm donor nuclei compared to IVF endoderm. **(b)** MA-plot comparing gene expression between endoderm samples in NT *versus* IVF-embryos. The mean log2-fold change (log2FC) gene expression in NT-embryos over IVF is plotted on the y-axis, while the mean log2(RPKM+1) gene expression in mesoderm donor nuclei is plotted on the x-axis. NT reprogramming from mesoderm to endoderm revealed 1737 ON-memory genes (purple) and 2013 OFF-memory genes (blue). **(c)** Boxplots showing log2(RPKM+1) mean expression levels of Reprogrammed-Down (RD; light gray), ON-memory (ON-mem; purple), Reprogrammed-Up (RU; dark gray), and OFF-memory (OFF-mem; blue) genes in mesoderm donor samples and endoderm IVF and NT samples. p-values for pairwise comparisons were calculated using Wilcoxon rank-sum test **(d)** Heatmap showing relative gene expression (z-score) of differentially expressed genes obtained by pairwise comparisons between mesoderm donor and IVF-endoderm, and between NT-endoderm and IVF-endoderm samples. Rows represent log2FC in expression levels over mean expression levels in IVF. Hierarchical clustering of rows classified these genes into four groups, note group 1 (ON-memory genes) and group 4 (OFF-memory genes). **(d)** Endoderm and meso-/ectoderm tissue was collected from neurula stage 21 embryos and fixed for ChIP-seq profiling of histone modifications (H3K4me1/me3, H3K9me3, H3K27ac/me3, H3K36me3, H3K79me3, H2AK119ub, H2A.X-f). **(f)** Identification of histone modifications enriched around the TSS or gene bodies on ON-memory or **(g)** OFF-memory genes in endoderm donor tissue or (h-i) meso-/ectoderm tissue. P-value in (f) and (g) * indicates a p-value <0.001, Kolmogorov-Smirnov-test.

Previous studies have established the importance of various histone modifications in establishing resistance to reprogramming by stabilizing OFF-memory^30–32^. In contrast, H3K4me3 was shown to act as a reprogramming barrier by stabilizing ON-memory^8^ in *Xenopus* and mammalian species^27–30,32^. To investigate if additional histone modifications act as reprogramming barriers by stabilizing ON-memory *in vivo*, we generated genome-wide profiles for a variety of histone modifications using ChIP-seq across meso-/ectoderm and endoderm donor tissues (**Figure 1e**). This included histone modifications associated with gene repression^33,34^: H3K9me3, H3K27me3 and H2AK119ub, marks associated with active gene expression^33,34^: H3K4me3, H3K27ac, H3K36me3 and H3K79me3, marks associated with active enhancers^35–38^: H3K27ac and H3K4me1, and H2A.xf1^39^ with a less well-documented role (**Figure S1 d,e,f**).

We initially addressed the levels of these chromatin marks around genomic regions of reprogramming resistant ON- and OFF-memory genes in their respective donor cell tissues (endoderm or meso-/ectoderm). Promoters of endoderm ON-memory genes show elevated H3K4me3 levels compared to reprogrammed-down genes (**Figure 1f**), as previously reported^8^. We additionally observed significant differences between endoderm ON-memory genes and reprogrammed-down genes in the levels of the active marks H3K27ac around promoter regions and H3K79me3 in the gene bodies (**Figure 1f)**. Histone modifications marking endoderm OFF-memory genes versus reprogrammed-up genes reveal no significant differences in the repressive marks H3K9me3, H3K27me3 and H2AK119ub in the endoderm donor cells used in this study (**Figure 1g**). It is possible, that these repressive marks are established during later differentiation stages of the endoderm tissue. A comparison of histone modification levels in meso-/ectoderm donor cells around the promoters or gene bodies (**Figure S1 d,e,f**), shows enrichment of H3K4me3, H3K27ac, and H3K79me3 for mesoderm ON-memory genes similar to what we observed for endoderm ON-memory genes (**Figure 1h**). Instead, mesoderm OFF-memory genes show significantly elevated H3K9me3 and H2AK119ub levels around their gene bodies (**Figure 1i**) compared to reprogrammed-up genes, suggesting that repressive chromatin states are established around OFF-memory genes in the mesoderm tissue. The other histone marks analyzed here did not reveal any significant differences around promoters or gene bodies of the ON- and OFF-memory gene sets (data not shown). Together, these analyses reveal that ON-memory correlates with specific histone modifications in the somatic cell donor nucleus and that this is consistent across tested cell types.

Taken together, transcriptional ON-memory seems to be a persistent biological feature of cellular reprogramming via NT mediated by similar chromatin features across different donor cell types. We furthermore generated reference datasets of cell-fate reprogramming suitable for building predictive computational models of reprogramming.

### Semi-supervised classification of memory genes reveals novel candidate reprogramming barriers marking ON-memory genes

Next, we asked which histone modifications, or combinations thereof, around the promoter regions of genes in the donor nuclei, are predictive of their memory class status after reprogramming in NT-embryos. In simple terms, we asked if using the knowledge about chromatin modifications around a gene in the somatic donor cell enables a prediction of whether this gene will be resistant or permissive to reprogramming in NT-embryos.

In an initial attempt to reveal potential predictive patterns in our combined chromatin and transcriptome profiling, we projected the memory class genes, along with their counterpart reprogrammed gene sets, onto a 2D PCA representation based on chromatin mark levels and gene expression patterns, with corresponding UMAP (**Figure 2 a,b**, **Figure S2 a,c)**. We note that in reduced dimensionality representations, ON- and OFF-memory genes are separated from one another. However, there remained overlap between ON-memory and reprogrammed-down (RD), and between OFF-memory and reprogrammed-up (RU) classes of genes. As memory class genes did not separate from reprogramming-susceptible genes in PCA, we hypothesized two possible scenarios: (1) gene memory status may exist on a continuum that cannot be unambiguously separated into discrete groups, or (2) it may be distinct combinations of histone modifications that confer an identifiable memory status.

**Figure 2:**
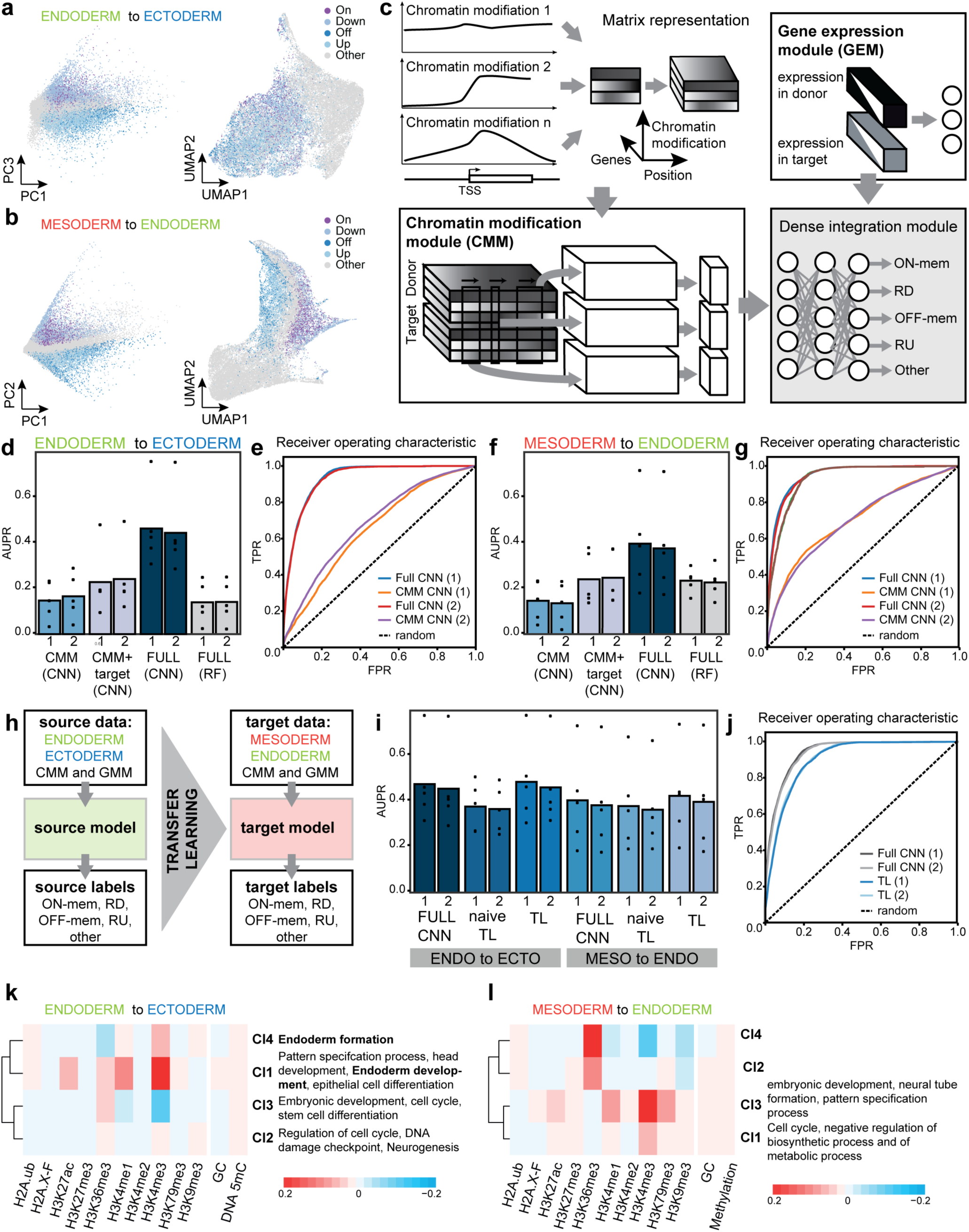
Digital Reprogramming predicts Memory genes in NT reprogramming and identifies novel epigenetic barriers. **(a-b)** PCA and UMAP representations of memory class genes visualized based on histone modification level and gene expression. **(c)** Schematic of Convolutional Neural Networks (CNNs) used to infer memory status. **(d)** Average area under Precision-Recall (AUPR) scores in endoderm to ectoderm reprogramming using different CNN and random forest (RF) models as indicated. **(d)** Receiver operating characteristic (ROC) curve for (d) endoderm to ectoderm reprogramming models as indicated, seed 1 and 2 **(f)** Average AUPR scores for somite to endoderm reprogramming. **(g)** ROC curve for (f) mesoderm to endoderm reprogramming models as indicated, seed 1 and 2 **(h)** Schematic for transfer learning used to gauge the effectiveness of learned features **(i)** Average PR scores in TL schemes. **(j)** ROC curves for (i). **(k and l**) Average activations of ON-memory genes over true positive genes were subclustered to identify different classes of ON-memory genes in endoderm to ectoderm (k) and mesoderm to endoderm reprogramming (l). Color indicates the sum over the cluster average activation of the 5kb TSS-flanking region, with higher scores indicating a greater contribution of that histone modification to predicted memory status.

To identify such potential hidden and more complex patterns in our data sets, we moved on to using machine learning algorithms to predict the reprogramming outcome of genes. Convolutional neural networks (CNNs) have been successfully used to make inferences from large-scale biological data, including methylation state from incomplete single-cell methylation data^40^, for predicting histone modifications from DNase and DNA-sequence data^41^, and for predicting gene expression from histone modifications^42^. Therefore, we tested next if CNN models are capable of inferring memory-class status based on a combination of gene expression and chromatin modifications around the promoter.

To this end, we first represented chromatin modification data in wild-type somatic donor cell types as a 1D array with 22 channels i.e., a one-dimension position along the transcriptional start site (TSS) with each histone modification representing an individual channel. We chose a window 5 kb up- and downstream of the TSS, so that potential instructive epigenetic information residing around promoter regions (around TSS), proximal enhancer regions (upstream of TSS), and gene body (downstream of TSS) were included. These so-generated channels were then combined in a “Chromatin Modification Module” (CMM; **Figure 2c**). Information on gene expression in the wild-type somatic donor cell type as well as on the wild-type target cell type (the cell type we aim at reprogramming to) was combined in a “Gene Expression Module” (GEM; **Figure 2c**). As output, we used the memory-class status (ON- or OFF memory; Reprogrammed-Up or -Down genes (**Figure 2c**). CNNs were then used to jointly integrate the so-represented histone modification data around the TSS and gene expression data (**Figure 2c** and Material and Methods). We then divided the input data into three parts: one to train the model, one for making predictions, and one for determining significance.

Using this processed input data, we then built three different CNN models to predict memory class status in the reprogrammed cell, each with increasing complexity to test how much information input is needed for successful prediction: The first model used only chromatin modification module as input to predict the memory class status in the reprogrammed cell (CMM-CNN). Second, a model used, in addition to the CMM, the gene expression data in the wild-type target somatic cell types (“<u>CMM</u>+target-CNN”). Third, the most complex model used the complete chromatin modification module and gene expression module in the donor and target cell type (FULL-CNN). To provide a baseline for the predictive performance of CNNs, we also used a random forest classification corresponding to the FULL-CNN using chromatin modification and gene expression in the donor and target cell type (FULL-RF).

Then, the prediction accuracy of each model was quantified using the Area Under Precision-Recall (AUPR; **Figure 2d,f**) or Receiver Operating Characteristics (ROC**; Figure S2b,d)** on a one-vs-all basis (**Figure 2 E,G and Figure S2 E,G**). All three CNNs could infer the memory status of a gene with accuracy comparable to or better than random forest classifiers, with both performing substantially better than expected by random classification (**Figure 2d-g**). Including both histone modification and gene expression information (FULL-CNN) in predictive models showed the greatest accuracy, with reduced models i.e., using donor tissue histone modification and target tissue expression (CMM+target CNN), still retaining accuracy (**Figure 2d-g**). Using histone modification alone provided the lowest classification accuracy, albeit also higher than random, with CNNs performing better than the corresponding RFs (**Figure 2d-g**). This suggests that chromatin modifications around genes hold information sufficient to determine whether a gene is likely to be an ON-memory gene or a reprogrammed-down gene, an OFF-memory or reprogrammed-up gene, or a gene that will not change its transcript expression levels during reprogramming. Information about the gene expression in the wild-type donor and target cell type, however, contains further information that can boost the prediction performance. In summary, our models’ predictive accuracy suggests that CNNs could identify useful predictive features from the provided data.

To investigate if our models were indeed learning useful histone modification combinations, we used transfer learning approaches to make predictions in NT reprogramming using alternative cell types. Specifically, we used the full TSS model trained on ectoderm-derived endoderm to make predictions about memory status in mesoderm-derived endoderm. We termed this naïve transfer learning (nTL), as CNNs were directly applied to new datasets with no additional parameter optimization (**Figure 2h**). A second form of transfer learning was also used, in which the network architecture and parameters were transferred, but with the parameters in the final dense layers of the network optimized using a subset of data (full transfer learning; TL; **Figure S2e**). We note that whilst the nTL showed a slight drop in performance compared to a fully tuned CNN, its performance remained better than random forest, and tuning of the densely connected layers showed performance comparable to that of a fully optimized network (**Figure 2h-j**). The observation that features in our CNNs could be transferred between cell types and still retain predictive performance suggests that useful representations of the features were indeed being learned in the upper layers of the network. Furthermore, it suggests that CNNs trained with one cell type can be used to predict the transcriptome reprogramming of any other cell type of interest.

Once we established the robust predictive accuracy of our model Digital Reprogramming, we turned to a second objective of our model, which was to identify possible classes of ON-memory genes regulated by different histone modifications or combinations thereof. To identify which of the histone modification features are predictive of memory status, we calculated the activations of the here-used genomic regions using DeepExplain^43,44^. Specifically, for the region 5 kb up- and downstream flanking the TSS of a given gene, we were able to calculate the activation contribution of a specific histone mark to a specific classifier e.g., the contribution of H3K4me3 in a given genomic interval around the TSS to ON-memory of that gene. To look for individual classes of memory status genes we calculated the overall activation contribution for each histone modification to ON-memory status and performed hierarchical clustering. We clustered the genes based on the similarity of the calculated activation contributions to individual subclasses (**Figure 2k**) and identified H3K4me3, H3K4me1, and H3K27ac as an important contributor to ON-memory status, but less so for reprogrammed-down status in both endoderm to ectoderm and mesoderm to endoderm reprogramming **(Figure 2g,h**).

Of the four subclusters formed by the endoderm ON-memory gene set, cluster 1 showed the strongest influence by H3K4me3, with cluster 4 and cluster 2 also showing some influence by H3K4me3. In addition to H3K4me3, we observed that cluster 1 also appeared to be influenced more strongly by H3K4me1 and H3K27ac than by other chromatin modifications, such as H3K36me3. Clusters 1, 2, and 4 were also influenced by DNA methylation and the GC content of the investigated gene region. Gene ontology enrichment analyses revealed that genes in clusters 1 and 4 were enriched for terms related to endoderm and other developmental signaling, indicating that this could be relevant chromatin modifications of genes important for donor cell identity. Cluster 3 was enriched for ontologies relating to the cell cycle, development and differentiation. Cluster 2, with no clear signature, was enriched for terms related to the cell cycle and DNA damage checkpoint.

A similar sub-clustering based on the activations of individual histone marks was done for mesoderm ON-memory genes (**Figure 2l**). H3K4me3 was again identified as an important contributor to ON-memory status, but not reprogrammed down status. Clusters 1 and 3 were influenced by H3K4me3, with cluster 3 being additionally influenced by H3K4me1, H3K27ac and H3K79me3. Cluster 2 and 4 were mainly influenced by H3K36me3. Gene ontology enrichment analyses (**Figure 2k**) revealed that genes in cluster 3 were associated with terms relating to development and pattern specification processes, and cluster 1 with cell cycle and metabolic processes. Cluster 2 and 4 showed no significant enrichment. Together, this suggests that clusters marked in both donor cell types by H3K4me1, H2K4me3, and H3K27ac are enriched for cell lineage genes. Such lineage genes have previously been determined to be ON-memory genes, whose correct reprogramming is essential for successful cell fate conversion in NT-embryos^5^. Therefore, H3K4me1, H3K4me3, and H3K27ac may represent an important combination of histone modifications stabilizing cell lineage genes and master ON-memory genes.

Together, our Digital Reprogramming approach accurately predicted transcriptional reprogramming outcomes by classifying genes as memory class genes or reprogrammed genes. Furthermore, Digital Reprogramming identified candidate reprogramming barriers linked to the inheritance of ON-memory, which were consistent between two donor cell types, endoderm and mesoderm. Finally, our model confirmed H3K4me3 as a key chromatin feature of ON-memory genes overall and identified H3K4me1 and H3K27ac as novel features.

### p300/CBP bromodomain inhibition in donor nuclei reduces ON-memory gene expression in NT-embryos

In addition to H3K4me3, we identified a predictive power of H3K4me1 and H3K27ac to infer the ON-memory state in our model and established a correlation between ON-memory gene expression. We could also observe H3K27ac enrichment on promoter regions of ON-memory genes when compared to reprogrammed-down genes. We are currently unable to interfere specifically with H3K4me1 due to technical limitations. Thus, we focused on H3K27ac and asked if its marking of promoter and/or enhancer regions contributes to the reprogramming resistance of ON-memory genes *in vivo*. To do so, we tested if reducing H3K27ac levels in donor nuclei could rescue the ON-memory phenotype in NT-embryos and improve nuclear reprogramming.

To perturb global H3K27ac levels in donor nuclei suitable for NT, we used SGC-CBP30 (hereinafter CBP30), a p300/CBP^45,46^ bromodomain inhibitor previously reported to selectively reduce H3K27ac levels ^47–49^. We treated embryos with CBP30 or DMSO as a control from the late gastrula stage until the neurula stage, i.e. 6 hours (**Figure 3a**). Western Blot confirmed the global reduction of H3K27ac levels in nuclei extracted from CBP30-treated embryos when compared to DMSO control (**Figure S3a**). To assess the effect of p300/CBP bromodomain inhibition on other histone modifications, we used liquid chromatography coupled to tandem mass-spectrometry (LC-MS/MS) to quantitatively assess post-translational modifications on histones isolated from CBP30- or DMSO-treated embryos. Analyzing lysine modifications on the H3 and H4 N-terminal tail revealed a mean 1.7-fold reduction (**Table S1**) of H3K27ac levels upon CBP30 treatment compared to DMSO, while H3K18ac levels remained constant in biological triplicates **(Figure 3b, Table S1**), in line with previous reports from cell culture systems^50^. Additionally, Western Blot analyses revealed a moderate reduction in H3K9ac levels upon CBP30 treatment, when compared to DMSO treatment (**Figure S3a**), potentially as a secondary effect of H3K27ac perturbation. Taken together, CBP30-inhibition of the p300/CBP-bromodomain led to globally reduced H3K27ac and H3K9ac levels in donor embryos.

**Figure 3:**
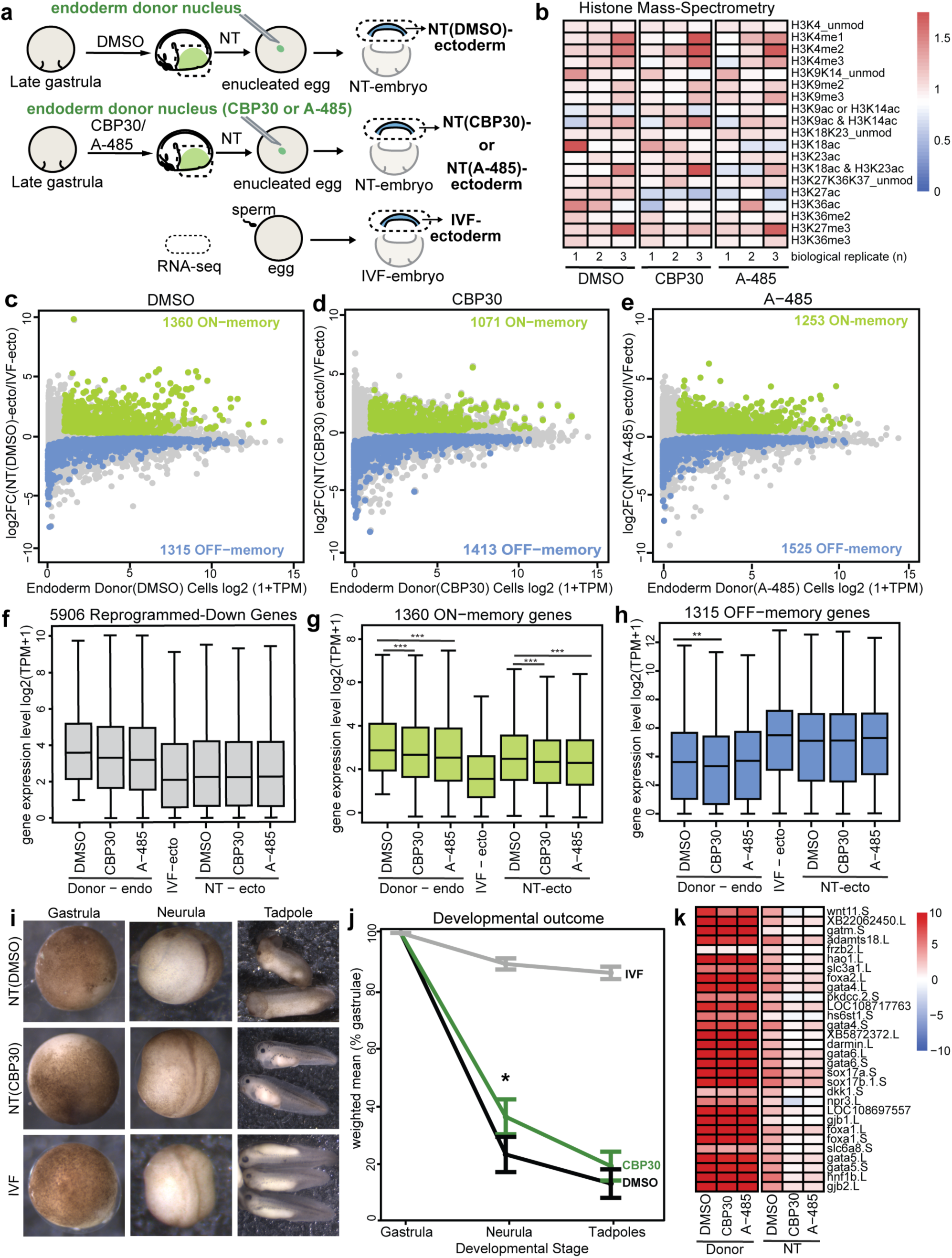
p300/CBP inhibition in endoderm donor nuclei reduces ON-memory gene expression in NT ectoderm samples. **(a)** Endoderm donor cell preparation under p300/CBP perturbed or control conditions, and NT experiments for endoderm to ectoderm reprogramming. **(b)** Histone mass-spectrometry assay. Heatmap representing candidate histone marks in DMSO, CBP30 and A-485 treated samples. Colors represent the relative histone mark abundance as fold-change over untreated samples, *n*=3 biological replicates. **(c-e)** CBP30 and A-485 treatment in donor nuclei reduces the number of ON-memory genes and increases the number of OFF-memory genes in NT-embryos. MA-plot comparing gene expression between (c) NT(DMSO) and IVF, (d) NT(CBP30) and IVF, and (d) NT(A-485) and IVF ectoderm samples. Mean log2FC in NT over IVF ectoderm expression levels is plotted on the y-axis, and the mean log2(TPM+1) expression in endoderm donor nuclei is plotted on the x-axis. Gray: all transcripts. Green: ON-memory genes. Blue: OFF-memory genes. **(f-g-h)** Boxplots comparing mean expression levels of (f) reprogrammed-down; (g) ON-memory and (h) OFF-memory transcripts in endoderm donor cells, IVF ectoderm, and NT ectoderm samples. Boxplots depict the median and the interquartile range (IQR). p-values for pairwise comparisons: Wilcoxon rank-sum test, two-sided (**p-values<0.01, ***p-values<0.001). **(i)** NT(DMSO), NT(CBP30) and IVF embryos at the gastrula, neurula and tadpole stage. **(j)** The development of NT(DMSO), NT(CBP30), and IVF gastrulae (st.10.5) was monitored until the tadpole stage. Y-axis represents the weighted mean percentage of gastrulating embryos reaching neurula (*p-value=0.02, one-tailed paired t-test) and tadpole stages (p-value=n.s.). **(k)** Heatmap depicting significantly downregulated genes in NT(CBP30) and NT(A-485) samples compared to NT(DMSO) (p-adj <0.05, log2 fold-change <0). Colors depict mean expression over IVF. Rows were sorted using hierarchical clustering, and Euclidean distance.

Next, we tested if p300/CBP bromodomain inhibition in donor nuclei can improve reprogramming in the resulting NT-embryos. For this, we injected CBP30-treated or DMSO-treated donor nuclei into enucleated eggs. As controls for a normal embryonic transcriptome, we *in vitro* fertilized eggs. We collected ectoderm tissue from gastrula stage NT(CBP30)-, NT(DMSO)- and IVF-embryos and subjected them to RNA-seq analysis along with endoderm donor(CBP30)- and donor(DMSO)-samples (**Figure 3a**). The experiments were performed in biological replicates as described in **Table S2**. Using differential gene expression analysis, we then compared transcriptomes between donor endoderm samples, corresponding NT-ectoderm samples, and IVF-ectoderm samples (**Figure S3c,d**, **Table S3**). We defined ON- and OFF-memory gene classes, as well as reprogrammed-down and -up genes as described above (**Table S4**).

This revealed that the number of ON-memory genes in NT(DMSO)-ectoderm samples was reduced from 1360 ON-memory genes to 1071 genes in NT(CBP30)-ectoderms (**Figure 3c-d**). When comparing the mean gene expression levels of ON-memory genes (**Figure 3f-g**), we find that it is moderately decreased in treated and control-treated donor tissues, and, importantly, decreased in NT(CBP30)-ectoderm tissues compared to NT(DMSO) ectoderm tissues. This indicated that p300/CBP bromodomain inhibition in donor nuclei can correct the ON-memory phenotype in NT-embryos. Interestingly, when we partitioned the ON-memory genes based on the fold-change in expression levels between NT(DMSO) *vs.* IVF or NT(CBP30) *vs.* IVF, we observed that the proportion of ON-memory genes with higher expression in NT(CBP30) *vs.* IVF was preferentially reduced compared to NT(DMSO) *vs.* IVF. In contrast, the proportion of ON-memory genes with the lowest NT *versus* IVF gene expression fold-change increased in NT(CBP30) samples (**Figure S3E**). This indicates, that CBP30-treatment in donor nuclei not only decreased the number of ON-memory genes (**Figure 3C,D**), but it also decreased the extent to which ON-memory genes are abnormally expressed in NT(CBP30) compared to IVF embryos.

On the other hand, we detected 1315 OFF-memory genes in NT(DMSO)-ectoderm samples and 1413 OFF-memory genes in NT(CBP30)-ectoderm samples (**Figure 3c,d**), suggesting an overall increase in the number of detected OFF-memory genes. However, the mean expression levels of these OFF-memory genes were similar in NT(CBP30) vs. NT(DMSO) samples (**Figure 3h**). Thus, our transcriptome assay suggests that CBP30 treatment in donor nuclei can improve ON-memory in NT-embryos, but not OFF-memory.

We then asked if the transcriptome in NT-embryos derived from CBP30-treated donor cells showed improved reprogramming and became more like that of IVF-embryos, when compared to the transcriptome of NT(DMSO)-embryos. PCA revealed that CBP30 and DMSO-treated donor endoderm samples grouped together, suggesting their transcriptomes are globally comparable (**Figure S3c**). Donor transcriptomes separated from all ectoderm samples along PC1. Along PC2, IVF-ectoderm transcriptomes separated from NT-ectoderm samples derived from both DMSO and CBP30-treated donor cells (**Figure S3c** We did not observe a closer association in PC space between NT(CBP30)-ectoderms and IVF-ectoderms than for NT(DMSO)-ectoderms and IVF-ectoderms (**Figure S3c-d**). This suggests that CBP30 treatment of the donor nuclei does not globally improve transcriptome reprogramming in the resulting NT-embryos.

Taken together, these experiments suggest that CBP30-mediated p300 inhibition in the donor cell correlates with a reduction in global H3K27ac levels and a loss of the ON-memory state of genes during nuclear reprogramming.

### p300/CBP HAT domain inhibition in donor nuclei reduces ON-memory gene expression in NT-embryos

Next, we tested if globally inhibiting the p300/CBP catalytic activity and thus reducing histone acetylation in donor cells mimics the effects of bromodomain inhibition on nuclear reprogramming. The p300/CBP bromodomain recognizes acetylated lysine residues on histones^49,51,52^, thus maintaining acetylation and recruiting transcriptional co-regulators. We hypothesized that targeting the p300/CBP histone acetyltransferase (HAT) domain, and not the architectural role of p300/CBP, could have distinct effects on transcriptional memory in reprogramming.

To address this, we used the small molecule A-485^53^, an acetyl-CoA competitive inhibitor, which prevents lysine acetylation on histones. We treated donor embryos with A-485, as described for CBP30 (**Figure 3a**). Compared to DMSO, A-485 treatment in donor embryos led to a 1.8-fold reduction in global H3K27ac levels, but also affected H3K18ac levels, as well as several other acetylated lysine residues on the histone H4 N-terminal tail (**Table S1**). Western Blot analysis also revealed reduced H3K9ac levels upon A-485 treatment (**Figure S3b**). In summary, A-485 treatment broadly affected histone H3 and H4 lysine acetylation in donor nuclei, while the CBP30 treatment depleted H3K27ac and H3K9ac.

We then tested the effects of HAT inhibition on reprogramming in NT-embryos. For this, we transplanted neurula stage donor cells treated with A-485 or DMSO to enucleated eggs and isolated ectoderm tissues of the resulting gastrula stage NT(A-485)- and NT(DMSO)-embryos (**Figure 3a** and **Table S2, S3 and S4**).

Differential gene expression analysis identified the different classes of reprogramming resistant and susceptible genes, as described above and in the methods section. This revealed 1253 ON-memory genes and 1525 OFF-memory genes in the ectoderm of NT(A-485)-embryos (**Figure 3e**). This represents a modest reduction in the number of ON-memory compared to the 1360 ON-memory genes identified in the ectoderm samples of NT(DMSO)-embryos (**Figure 3c**). We observe an increase in the number of OFF-memory genes in NT(A485)-embryos when compared to NT(DMSO)-ectoderm from 1315 to 1252 (**Figure 3c,e**). Therefore, our findings indicate that p300/CBP HAT inhibition reduces the number of ON-memory genes but increases the number of OFF-memory genes in NT-embryos, similar to p300/CBP bromodomain inhibition. The mean expression levels of ON-memory genes decreased in NT(A-485)-ectoderm, when compared to NT(DMSO), as well as in the treated donor tissue compared to the untreated one. The mean expression levels of OFF-memory genes did not significantly increase and remained comparable between NT(A-485)-ectoderm and NT(DMSO) (**Figure 3g,h**). Finally, PCA analyses did not reveal a global transcriptome correction in NT(A-485) ectoderm tissues, as their transcriptomes clustered together with NT(DMSO) samples and separated from IVF-samples (**Figure S3c**).

Together, this suggests that p300/CBP HAT inhibition results in a modest improvement in reprogramming ON-memory genes in NT-embryos.

### p300/CBP bromodomain inhibition moderately improves the developmental outcome of cloned embryos

Since we observed an improvement of ON-memory upon p300/CBP inhibition, albeit modest, and an associated reduction in H3K27ac levels, we asked next if this also correlates with improved cell fate reprogramming. Therefore, we perturbed p300/CBP-mediated histone acetylation using CBP30 inhibitors to reduce ON-memory in NT-embryos and tested the effect on the developmental outcome of NT-embryos.

To this end, we selected the bromodomain inhibition approach to perturb the donor nuclei before the developmental assay, since CBP30 treatment mainly affected H3K27ac levels in donor nuclei and only rescued ON-memory in NT-embryos, leaving OFF-memory unperturbed. Hence, we generated cloned embryos from DMSO and CBP30-treated donor nuclei. Both sets of NT-embryos demonstrated similar developmental rates during the blastula and early gastrula stage, but we observed healthier morphology in NT(CBP30) embryos, which reflected higher rates of development compared to NT(DMSO) embryos at the neurula stage (**Figure 3 i,j, Figure S3h, Table S5**).

We have previously reported that endoderm master regulators display ON-memory when endoderm is reprogrammed to ectoderm fate in NT-embryos and that their aberrant expression causes differentiation defects in NT-ectoderm^5,6^. Since we observed improved development in NT(CBP30)-embryos, we speculated whether this is due to a correction of this subset of ON-memory genes.

Thus, we tested if p300/CBP inhibitor treatment in donor nuclei can reduce the ON-memory expression of endoderm marker genes in NT-ectoderm and found that lineage transcription factors *gata6*, *sox17a*, and *sox17b*, as well as endoderm genes such as *darmin*, *wnt11*, and others were expressed at significantly lower levels in NT(CBP30) and NT(A-485) samples compared to NT(DMSO), despite retaining high expression levels in p300/CBP-perturbed donor samples (**Figure 3k**). Thus, this set of genes became permissive to reprogramming upon p300/CBP perturbation, which we termed ‘treatment-sensitive’ genes. On the other hand, the endoderm lineage TF *foxa4* and the endoderm marker gene *a2m* retained high ON-memory expression in NT(CBP30) and NT(A-485) despite p300/CBP perturbation in donor nuclei reprogramming (**Figure S3g**), thus being ‘treatment-insensitive’. Therefore, we hypothesized that the p300/CBP inhibitor approaches may have reduced H3K27ac levels on the promoters of treatment-sensitive, but not on insensitive genes, explaining their different behavior upon nuclear reprogramming. Surprisingly, using ChIP-qPCR in donor nuclei, we found reduced H3K27ac levels around the promoters of both sensitive (such as *sox17a* and *gata5*) and insensitive genes (such as *a2m* and *foxa4*) upon CBP30 treatment (**Figure S3 f,g**). This suggested that reduced H3K27ac levels around ON-memory gene promoters alone were insufficient to explain whether genes would lose their ON-memory status in NT embryos upon CBP30 treatment in donor nuclei.

Thus, our findings suggest that p300/CBP inhibition reduces transcriptional ON-memory of a key set of endoderm genes, which correlates with improved NT-embryo development. This indicates that p300/CBP and its unperturbed activity contribute to the barriers of cell fate reprogramming and to the mechanisms safeguarding cellular identities.

### Proximity to active enhancers distinguishes ON-memory genes from correctly reprogrammed genes

Having established a role for p300/CBP and associated H3K27ac as a barrier for reprogramming, we aimed to discern the mechanisms underlying H3K27ac-mediated ON-memory in more detail. We asked if H3K27ac around enhancers or promoters, or both, could participate in maintaining ON-memory in nuclear reprogramming. For this, we first correlated H3K27ac levels around enhancers in donor cells with ON-memory status in NT reprogramming.

To identify putative enhancer elements in endoderm donor nuclei, we analyzed a previously published ChIP-seq dataset^54^ addressing p300 binding in endoderm tissue from wildtype neurula-stage embryos. We classified the peaks into promoter peaks (TSS +/-500 bp) and non-promoter peaks (at least 500 bp away from the promoter). Next, we selected the non-promoter p300 peaks as a proxy for putative enhancer sites. We assigned the p300 peaks to their putative target genes^55^ based on the following proximity criteria: (1) p300-peaks overlapping a gene body were assigned to that gene, (2) p300-peaks overlapping more than one gene or not overlapping any genes were assigned to the nearest promoter, (3) the target gene is actively expressed in endoderm tissue at the neurula stage (TPM>1). For simplicity, we will further refer to the identified non-promoter p300-peaks proximal to genes as “enhancers of those genes”. We next compared the enrichment levels for H3K27ac on enhancers proximal to ON-memory and reprogrammed-down genes (**Figure 4a**) and observed a significantly higher H3K27ac signal for ON-memory compared to RD genes. This suggests that, similar to promoters (**Figure 1f**), putative enhancers of ON-memory genes are enriched for H3K27ac compared to properly reprogrammed-down genes.

**Figure 4:**
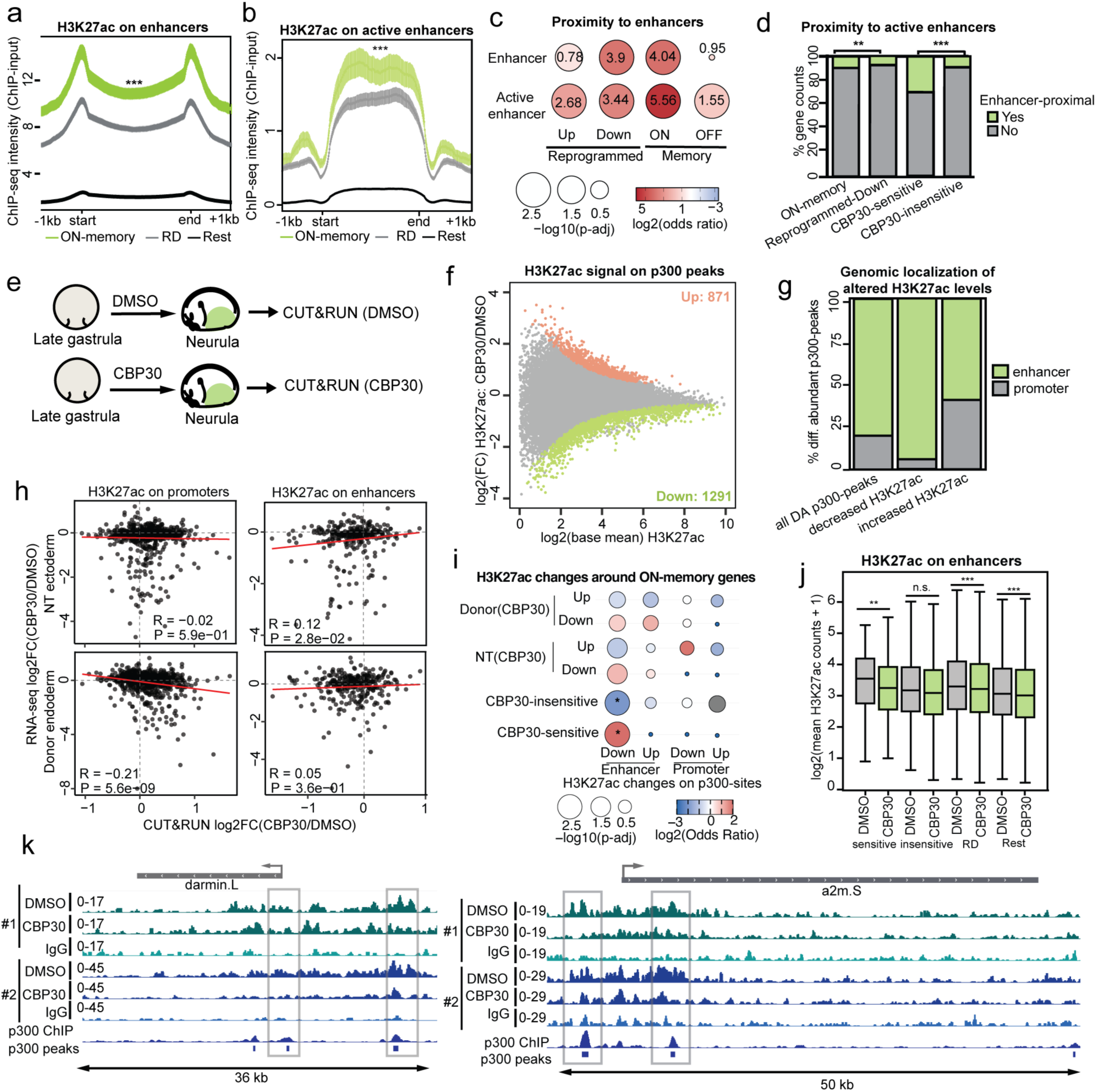
ON-memory genes are enriched for active enhancers, which are targeted by CBP30 and demonstrate reduced H3K27ac levels. **(a-b)** H3K27ac ChIP-seq intensity around (a) p300-sites +/-1 kb at predicted enhancers and **(b)** predicted active enhancers in neurula-stage endoderm. ChIP-seq intensities are higher for enhancers and active enhancers proximal to ON-memory genes (green) *vs.* RD genes (gray). (For **(a)** enhancers: p-value <10^-24^ (***), and **(b)** active enhancers p-value<10^-3^ (***), Wilcoxon rank-sum test). Genes excluding ON-memory and RD genes (rest): black. **(c)** Balloon plot comparing enrichment of enhancer-proximal reprogramming genes *vs.* all genes in endoderm nuclei. Balloon sizes represent p-adj; colors represent the log2 odds ratio. Balloons are labeled with the raw odds ratio of each comparison. **(d)** Proportions of ON-memory and RD (p-value = 2·10^-5^) genes proximal (green) or not proximal (gray) to an active enhancer (green), and between CBP30-sensitive (n=36 genes) and CBP30-insensitive genes (n=1324 genes; p-value<10^-3^). **(d)** CUT&RUN in donor embryos treated with DMSO or CBP30. Endoderm tissues (green) were isolated for CUT&RUN against H3K27ac. **(f)** MA-plot comparing H3K27ac levels in donor(CBP30) vs. donor(DMSO) endoderm on p300-peaks. Y-axis: mean log2FC (CBP30/DMSO); x-axis: base mean H3K27ac. Gray: all p300-peaks; green: p300-peaks with decreased H3K27ac levels (p-value <0.05), orange: p300-peaks with increased H3K27ac levels (p-value <0.05). **(g)** Genomic distribution of 2162 differentially abundant (DA) p300-peaks with H3K27ac, of which 1291 DA p300-peaks with decreased and 871 p300-peaks with increased H3K27ac levels on enhancers (green) or promoters (gray). **(h)** Scatter plot comparing the log2FC in ON-memory gene expression for NT(CBP30/DMSO) or donor(CBP30/DMSO), against log2FC in H3K27ac signal for donor(CBP30/DMSO) on promoters and active enhancers. Pearson’s correlation coefficients and p-values are shown within each box. **(i)** Balloon plot comparing subsets of ON-memory genes based on gene expression changes in donor(CBP30/DMSO), or in NT(CBP30/DMSO), against changes in H3K27ac levels on promoters or enhancers. Balloon sizes: -log10 p-adj; colors: log2 odds ratio. **(j)** Boxplot comparing H3K27ac levels in donor(DMSO) and donor(CBP30) on enhancers proximal to treatment-sensitive ON-memory genes (n=157 peaks, p-value = 0.03), treatment-insensitive ON-memory genes (n=4647, p-value = 10^-5^), RD genes (n=14841, p-value =1.09·10^-9^), and Rest (n=33256, p-value = 3.74·10^-16^). Wilcoxon rank-sum test. **(k)** Genome browser snapshots for the ON-memory genes *darmin.L* and *a2m.S*.

As H3K27ac has been reported to mark active enhancers^35–38,56^, we tested if the co-occupancy of p300- and H3K27ac-peaks outside of promoters, marking a putative active enhancer, correlates with maintaining an ON-memory state from endoderm donor nuclei to the ectoderm of NT-embryos. To this end, we called broad H3K27ac peaks in ChIP-seq data from wildtype endoderm nuclei (**Figure 1f, Figure S1d, Figure S4a**) and intersected the H3K27ac-peak set with the p300-peak set described above to obtain a set of H3K27ac+/p300+ putative active enhancers. This prediction of an active enhancer set was supported by the presence of detectable H3K4me1 signal on these sites (**Figure S4a**), as well as undetectable RNA-seq transcript signal from the non-promoter sequences (data not shown). We thus curated a list of 1306 putative active enhancers based on H3K27ac and p300-peak overlap and paired them with 1066 genes, fulfilling the same criteria described for the p300-peak set above. Resembling our findings around enhancers overall (**Figure 4a**), we observed a strongly enriched H3K27ac signal around active enhancers (**Figure 4b**) for ON-memory genes compared to reprogrammed-down genes in the somatic donor cell.

Interestingly, when we compared the genes paired to a putative enhancer only marked by a p300-peak, and the groups of memory and reprogrammed genes, we detected significant enrichment for both ON- and OFF-memory genes, as well as RD and RU genes, albeit the highest odds ratios were detected for ON-memory genes (**Figure 4c**). Using the set of putative active enhancers, co-occupied by p300 and H3K27ac, we observed enrichment only for ON-memory and reprogrammed-down genes compared to all genes (**Figure 4c**), in line with the active expression of these two gene sets in endoderm cells of neurula-stage embryos. OFF-memory and reprogrammed-up genes, which are lowly expressed or undetectable in endoderm cells at this stage, were not enriched for genes proximal to putative active enhancers **(Figure 4c)**. Excitingly, when comparing ON-memory and reprogrammed-down genes, the group of ON-memory genes was more enriched for enhancer-proximal genes than reprogrammed-down genes (**Figure 4d**). This hinted that the proximity to an H3K27ac-marked p300-site outside of the promoter region, i.e., an active enhancer, could be correlated to the memory status of ON-memory genes.

Then, we tested if the proximity to an active enhancer correlates with the treatment sensitivity of ON-memory genes to CBP30 or A-485. Indeed, we found that 30% of the CBP30-sensitive ON-memory genes of the CBP30-sensitive ON-memory genes were proximal to an active enhancer, in contrast to 9,2% of the CBP30-insensitive ON-memory genes (**Figure 4d**).

These findings hint that proximity to active enhancers may correlate with ON-memory gene expression sensitivity to CBP30 treatment and loss of ON-memory status upon NT-reprogramming.

### Reduction of H3K27ac levels around enhancers of ON-memory genes correlates with improved reprogramming after NT

Having established increased H3K27ac marking of enhancers and promoters of ON-memory genes, we asked next if p300/CBP bromodomain inhibition, implicated in enhancer activity^51^, particularly affected H3K27ac levels around active enhancers, rather than promoters, in endoderm donor nuclei. Thus, we tested if H3K27ac levels changed in a genome-wide localized manner upon treatment with p300/CBP inhibitors in donor nuclei, and if this correlates with the treatment sensitivity of ON-memory genes in nuclear reprogramming.

To address this question, we performed a Cleavage Under Targets and Release Using Nuclease (CUT&RUN) assay against H3K27ac in endoderm donor nuclei treated with DMSO or CBP30 (**Figure 4e**). To compare the H3K27ac levels in DMSO control *vs.* CBP30 treated samples (**Figure S4b**), we used the p300-peak set described above as a reference peak set for sampling H3K27ac CUT&RUN reads for each condition and performed differential peak analysis. PCA of spike-in normalized H3K27ac counts around p300-sites revealed a clear separation between DMSO and CBP30 samples along PC1 (**Figure S4c**). Pairwise comparison between DMSO and CBP30 samples revealed 1291 differentially abundant p300-peaks (p-value<0.05), for which H3K27ac levels were decreased upon CBP30-treatment, and 871 peaks for which H3K27ac levels were increased (**Figure 4f**). Most of the differentially abundant p300-peaks were located outside of promoters (**Figure 4g**). Interestingly, when we divided the differentially abundant p300-peaks into peaks with decreased H3K27ac or increased H3K27ac levels upon CBP30-treatment, we observed that 94,3% of the p300-peaks with decreased H3K27ac levels were located on putative enhancers defined by non-promoter p300-peaks, while 40,6% of the p300-peaks with increased H3K27ac levels were located around promoters (**Figure 4g**). This suggests that CBP30 treatment mainly reduced H3K27ac on enhancers.

To further understand the correlation between changes in H3K27ac on gene regulatory elements in somatic donor nuclei and the reduced ON-memory expression in NT-embryos, we focused our analysis on gene regulatory elements paired with ON-memory genes. To this end, we performed correlation analyses comparing the ON-memory gene expression changes between NT(CBP30) and NT(DMSO) and the changes in H3K27ac between donor(CBP30) and donor(DMSO). Under CBP30 conditions, changes in H3K27ac levels on promoters in donors did not correlate with ON-memory gene expression changes in NT(CBP30) embryos compared to NT(DMSO). On the other hand, we detected a positive albeit weak correlation between CBP30-induced H3K27ac reduction on active enhancers (union of H3K27ac-peaks called in at least three biological replicates per condition in the CUT&RUN dataset, and p300-peaks) and reduced ON-memory expression in NT(CBP30) expression (**Figure 4h**). In contrast, changes in H3K27ac on either promoters or enhancers did not reveal any significant correlation with gene expression changes in donor(CBP30) nuclei (**Figure 4h**). Thus, we observe that perturbing H3K27ac levels in donor nuclei does not correlate with altered ON-memory gene expression in donor(CBP30) cells, but rather correlates with gene expression changes upon reprogramming, detected in NT(CBP30) embryos.

Having observed a correlation between CBP30-induced H3K27ac alterations on enhancers and gene expression changes in NT(CBP30) embryos, we asked if distinct groups of ON-memory genes displayed distinct changes in H3K27ac levels (**Figure 4i, S4e**). Thus, we stratified ON-memory genes based on their gene expression changes in NT(CBP30) vs. NT(DMSO) embryos and donor(CBP30) vs. donor(DMSO) embryos. Enrichment analysis revealed a significant over-representation of enhancers with decreased H3K27ac levels within the ON-memory gene set that was downregulated in NT(CBP30) *vs.* NT(DMSO) embryos (log2FC < −1). We then performed these enrichment analyses comparing enhancers with reduced H3K27ac levels to differentially expressed genes in donor samples (CBP30 *vs*. DMSO). We did not detect any significant enrichment, suggesting that, upon CBP30 treatment, changes in H3K27ac levels around enhancers of genes in donor cells are not indicative of changes in their expression levels before reprogramming. Instead, the CBP30-induced decrease in H3K27ac levels around enhancers is indicative of gene expression changes in their target genes only after reprogramming in NT-embryos.

When we focused on the set of ‘CBP30-sensitive’ genes (NT(CBP30) *vs.* NT(DMSO) p-adj<0.05, log2FC <0), we found the strongest enrichment for genes paired to enhancers with reduced H3K27ac levels upon CBP30-treatment in donor nuclei (**Figure 4i**). This led us to hypothesize that the reduction of H3K27ac levels on putative enhancers via p300/CBP bromodomain inhibition could underlie the loss of ON-memory status in nuclear reprogramming.

Finally, we compared the H3K27ac levels on CBP30-sensitive and insensitive genes between DMSO and CBP30-treated samples, focusing on putative enhancers. We observed a significantly reduced H3K27ac signal in CBP30 samples on enhancers associated with ‘CBP30-sensitive’ ON-memory genes. While H3K27ac levels also moderately decreased on enhancers associated with ‘CBP30-insensitive’ genes, this difference was non-significant. We also observed a mild decrease in H3K27ac levels on enhancers associated with reprogrammed-down genes or other genes, yet to a lower extent than for the set of peaks proximal to ‘CBP30-sensitive’ genes (**Figure 4 j,k, Figure S4d**). Together, this suggests that CBP30 treatment reduces H3K27ac levels around putative enhancers of a set of ON-memory genes and renders them permissive to reprogramming.

In summary, our analyses demonstrate that p300/CBP bromodomain inhibition leads to reduced H3K27ac levels at enhancers near ON-memory genes in donor nuclei, and these chromatin changes correlate with decreased ON-memory gene expression after nuclear transfer. This points towards a new role for H3K27ac on enhancers and the p300/CBP bromodomain in maintaining and reprogramming cell fates via somatic cell nuclear transfer.

## DISCUSSION

In the present study, we developed Digital Reprogramming, a convolutional neural network model that accurately predicts reprogramming outcomes in nuclear reprogramming by combining chromatin features and transcriptional profiles. Crucially, our “digital reprogramming” was also able to identify robust histone features responsible for memory status. Identified features included the known reprogramming barrier H3K4me3, a key epigenetic feature preventing cell fate changes, and a novel candidate barrier for nuclear reprogramming in NT-embryos, H3K27ac. H3K27ac was enriched around the promoters and putative enhancers of ON-memory genes. Excitingly, perturbing H3K27ac in donor nuclei via p300/CBP inhibition reduced ON-memory gene expression in the resulting NT-embryos and improved the developmental outcome of the cloned embryos. These results link p300/CBP activity and H3K27ac to the memory of active gene expression states during nuclear reprogramming and indicate them as safeguarding mechanisms of cellular identities.

Our model leveraged a large-scale reference dataset containing chromatin and gene expression modules to predict reprogramming resistance in *Xenopus* SCNT. Interestingly, we observed that the epigenetic features identified as novel reprogramming barriers, such as H3K27ac, had a consistent enrichment pattern on memory class genes compared to correctly reprogrammed genes in both donor cell lineages. This is an important hint, that maintaining an active chromatin state not only plays a role in maintaining cell fates, but it also requires accurate reprogramming to ensure transcriptome fidelity in the target cell type in NT-embryos.

The fact that transfer learning could be used to predict reprogramming efficiency of other cell types that were not used to train the model, suggests that Digital Reprogramming may be used to predict reprogramming efficiency also in human reprogramming systems. In this way, important reprogramming resistant cell lineage genes can be identified *in silico*, alongside the associated histone modifications acting as barriers. Interreference strategies can be designed ahead of the actual experiment, thus saving time and improving current reprogramming protocols.

Indeed, transcription factor (TF)-reprogramming, using a p300/CBP bromodomain inhibitor to reduce H3K27ac levels in fibroblasts improved iPSC reprogramming by decreasing the expression of fibroblast-specific genes when applied in the early stages of reprogramming^16^. Applying these inhibitors during iPSC reprogramming did not affect the activation of pluripotency gene expression. In our experimental setup, short-term p300/CBP inhibitor treatment was sufficient to deplete H3K27ac levels in donor cells and reduce ON-memory in the resulting, unperturbed, NT-embryos. Using histone mass-spectrometry analyses of the donor cell chromatin, we confirmed that p300/CBP bromodomain inhibition predominantly affected H3K27ac levels, while catalytic inhibition perturbed histone acetylation broadly in donor nuclei, yet remarkably both approaches corrected ON-memory in the resulting unperturbed NT-embryos similarly.

A key difference between NT in *Xenopus* and iPSC reprogramming is that TF-mediated reprogramming is accompanied by ongoing transcription, while early *Xenopus* embryos undergo 12 cycles of rapid cell divisions in a transcription-free window before zygotic genome activation (ZGA)^57,58^. In TF-mediated reprogramming, inactivating the transcriptional program of the starting cell type and establishing the transcriptional program of the target cell type occur hand in hand, thus complicating the distinction of whether perturbing a chromatin feature improves reprogramming due to diminishing the cellular memory status of the somatic cell or improving the establishment of the target transcriptome. In our model system, the perturbation of p300/CBP and H3K27ac in the donor cell is achieved first. Then, using NT, a zygote with unperturbed p300/CBP activities is formed, and the cloned zygote divides several times in the absence of transcription until zygotic genome activation (ZGA), when transcription occurs again^57,58^. Thus, experiments using this model system point towards a role for p300/CBP and H3K27ac in maintaining transcriptional ON-memory from the donor nucleus to the reprogrammed cell type during reprogramming.

Treatments of NT-embryos and iPSCs with histone deacetylase (HDAC) inhibitors have been previously used to improve reprogramming by increasing pluripotency gene activation ^59–64,65–69^. It has therefore been suggested that high levels of H3K27ac are beneficial for nuclear reprogramming. This is not in conflict with our observation. We propose that reducing H3K27ac only in the donor cell, leaving the histone acetyltransferase activities unperturbed after NT, allows successful inactivation of ON-memory genes in the resulting NT-embryos. A subsequent treatment of NT-embryos with HDAC inhibitors may then improve the activation of OFF-memory genes during reprogramming.

We detected significant H3K27ac enrichment on ON-memory gene promoters and proximal p300-sites or putative enhancers linked to ON-memory genes. Accordingly, we found that the ON-memory genes that were readily reprogrammed via SCNT upon p300/CBP bromodomain perturbation in the donor nuclei showed significant enrichment for proximity to an enhancer. Moreover, genome-wide analysis using CUT&RUN revealed depleted H3K27ac on enhancers coupled to ON-memory genes sensitive to p300/CBP bromodomain inhibition, and a positive correlation between H3K27ac levels on active enhancers, and ON-memory gene expression in NT embryos. While our analysis could not account for distal enhancer regulation of ON-memory genes, we detected a link between proximal H3K27ac-marked active enhancers and ON-memory.

While perturbing H3K27ac levels on enhancers led to improvements in the NT-embryo transcriptome, and moderately improved the development of NT embryos derived from p300/CBP bromodomain inhibitor-treated donor cells, there remained ON-memory expression in these cloned embryos. We speculate, that unperturbed H3K4me3 levels on ON-memory gene promoters could continue to maintain ON-memory expression, despite perturbed H3K27ac levels. Thus, this may hint that H3K4me3 on promoters, supported by H3K27ac on enhancers could act in concert to maintain an active expression state from the donor nucleus to the cloned embryo. However, further experiments involving genome region-specific manipulation of H3K27ac and H3K4me3, e.g. via dCas9-mediated targeting of chromatin modifiers, are required to test this.

Recent studies^70,71^ have challenged the previously attributed role^35,36^ of H3K27ac on active enhancers. Using histone mutagenesis, Sankar *et al.*^71^ observed minimal effects on active transcription under steady states, but only upon challenging cell identities by inducing differentiation. In our system, where briefly targeting the p300/CBP bromodomain mostly led to decreased H3K27ac levels on enhancers, we found that gene expression in donor nuclei did not correlate with changes in H3K27ac levels. Only after SCNT, when we challenged the steady-state by inducing nuclear reprogramming, we detected a correlation between gene expression in NT-ectoderm and H3K27ac changes in the donor cell. While our approach chemically targets p300/CBP, we cannot rule out effects due to altered chromatin accessibility or changes in the nuclear acetylome. However, the brief treatment window and our histone mass-spectrometry data showing high selectivity for H3K27ac help minimize such off-target effects. Therefore, our results align with findings that H3K27ac perturbation has minimal impact under steady-state conditions while suggesting that H3K27ac maintains cell state stability and, when removed, introduces a vulnerability for cell fates that could be exploited for cellular reprogramming.

Given the rapid turnover and dynamic nature of histone acetylation, the mechanism of how H3K27ac could contribute to ON-memory propagation during cellular reprogramming remains to be addressed. This implies that there could be a transcription-independent mechanism to maintain histone acetylation across mitosis and instruct aberrant memory gene expression in our model. In the fly embryo, Samata *et al.*^72^ recently reported that H4K16ac maintained from the oocyte to the embryo was retained on gene promoters preceding ZGA and instructed future gene expression via MOF retention on chromatin throughout mitosis. While this has not yet been shown for H3K27ac, Wong *et al*.^73^ have suggested that p300 is retained on promoters during mitosis, thus maintaining transcriptional memory. While the transcription-free window between fertilization and ZGA in *Xenopus* is poorly described, reports from previous species suggest that H3K27ac and p300/CBP are important for activating zygotic transcription^74–77^. Whether p300/CBP or H3K27ac arise *de novo* in the embryo or are maintained from the gametes is unclear, but our findings that H3K27ac from the donor nucleus maintains transcriptional memory in NT reprogramming provoke further mechanistic investigation in the wildtype embryo as well.

In summary, the Digital Reprogramming model presented here, successfully identifies barriers consistent across different cell types in donor nuclei via transfer learning, suggesting a persistent and robust biological feature. This offers many possibilities for its broader application across species and in various reprogramming contexts. The ability of our model to learn useful combinations of chromatin features and transcriptional patterns makes it a suitable platform for further training and development. In particular, incorporating information about TF binding, investigating histone modification patterns on enhancers, or including data acquired from single-cell approaches could pinpoint crucial cell fate vulnerabilities to be exploited in reprogramming. Further refinement of this model could provide accurate predictions and *in vivo* testable hypotheses that deepen our understanding of cellular identity, make accurate predictions of histone state in novel conditions, and accelerate discoveries in regenerative medicine.

## MATERIALS AND METHODS

### Xenopus Laevis culture

Adult *Xenopus Laevis* were obtained from Nasco (901 Janesville Avenue, P.O. Box 901, Fort Atkinson, WI 53538-0901, USA) and Xenopus1 (Xenopus1, Corp. 5654 Merkel Rd. Dexter, MI. 48130). All frog maintenance and care were conducted according to the German Animal Welfare Act. Research animals were used following guidelines approved and licensed by ROB-55.2-2532.Vet_02-23-126.

### *In vitro* fertilization

Eggs were *in vitro* fertilized by mixing with a sperm slurry in a Petri dish, incubating for 2 min at RT and flooding with distilled water, then dejellied using 2% cysteine solution pH 7.8 (adjusted using NaOH), washed 3 times with 0.1x MMR (100 mM NaCl, 2 mM KCl, 1 mM MgSO4, 2 mM CaCl2, 0.1 mM EDTA, 5 mM HEPES (pH 7.8)) and transferred into 0.1x MMR for culturing until the desired stage for endoderm donor cell preparation.

### Donor cell preparation for nuclear transfer (NT)

Endoderm and mesoderm were dissected from neurula-stage embryos (NF stage 18) and allowed to dissociate into single cells in calcium- and magnesium-free 1x modified Barth saline (MBS, 88 mM NaCl, 1mM KCl, 10mM HEPES, 2.5 mM NaHCO_3_, pH 7.4) with 1 mM EDTA and 0.1% bovine serum albumin (BSA) in a Petri dish coated with 0.1% agarose in H_2_O. Donor cells were immediately used for NT.

### Nuclear transfer (NT) and embryo culture

Nuclear transfer was performed as described previously^3^. Endoderm or mesoderm donor cells were partly disrupted by suction into a glass capillary needle and injected into an egg enucleated by UV exposure in a UV-crosslinker (CL-1000 Analytik Jena, exposure setting 2000). Nuclear transfer was performed immediately upon enucleating the acceptor eggs. The injected eggs were then placed in 1x MMR and the medium was changed to 0.1x MMR after cleavage of the embryos. Alongside, IVFs were cultured as control overnight at 16°C until they reached the blastula stage. Then, NT-embryos morphologically indistinguishable from IVF controls were selected for subsequent experiments. For gene expression analyses in gastrulae, NT- and IVF-embryos with the same blastopore size were selected to ensure equal developmental stages. The embryonic non-neural ectoderm or endoderm was excised from each embryo and snap-frozen on dry ice for RNA isolation^6^. To score the developmental outcome, the NT- and IVF-embryos were cultured in 0.1x MMR at 16°C until they reached the neurula stage and then at 23°C until the feeding tadpole stage.

### Total RNA extraction

Embryonic tissues were collected and extracted as described above and stored at −80°C. Total RNA was isolated using the RNeasy Mini Kit (QIAGEN, 74104) according to the manufacturer’s instructions. To lyse the embryonic tissue, samples were vortexed at high speed for 10 min at 4°C. DNase digestion was performed according to the manufacturer’s instructions, using RNase-free DNase (QIAGEN, 79254). RNA was eluted in 35 µl nuclease-free H_2_O.

### mRNA sequencing library preparation

Total RNA was quantified on a Qubit fluorometer, and the sample quality was assessed on a TapeStation before library preparation. Per sample, 400 ng total RNA of animal cap (ectoderm) tissue and 300 ng of endoderm donor tissue were used to isolate mRNA using the NEBNext Poly(A) mRNA Magnetic Isolation Module (NEB, E7490). Sequencing libraries were generated using the NEBNext Ultra II Directional RNA Library Prep Kit for Illumina (NEB, E7760), following the manufacturer’s instructions, using 12-13 PCR amplification cycles. Libraries were multiplexed and sequenced on a NextSeq 6000 platform.

### Chromatin Immunoprecipitation (ChIP)

Chromatin Immunoprecipitation (ChIP) was performed as described previously^8^. Briefly, for each ChIP experiment, 75 embryos were dissected in 1x MBS to divide the endoderm from the mesectoderm tissue. Samples were fixed in 1% formaldehyde in 0.1x MMR for 25 min at RT, washed with 0.1x MMR, equilibrated in 500 μL HEG solution (50 mM HEPES-KOH pH 7.5, 1 mM EDTA, 20% glycerol) and frozen at −80°C. To extract chromatin, the samples were homogenized in E1 (50 mM HEPES-KOH pH 7.5, 140 mM NaCl, 1 mM EDTA pH 8.0, 10% glycerol, 0.5% Igepal CA-630, 0.25% Triton X-100, 1 mM DTT, complete protease inhibitors (Roche)), washed with buffer E1, buffer E2 (10 mM Tris pH 8.0, 200 mM NaCl, 1 mM EDTA, 0.5 mM EGTA, complete protease inhibitors (Roche)) and buffer E3 (10 mM Tris pH 8.0, 200 mM NaCl, 1 mM EDTA, 0.5 mM EGTA, 0.1% Na-deoxycholate, 0.5% Na-laoroylsarcosine, complete protease inhibitors (Roche)), then resuspended in buffer E3. Chromatin was fragmented by sonication for 20 cycles (30” on and 30” off) using a Bioruptor (Diagenode) at 4°C, then centrifuged for 15 min at 4°C at full speed. The supernatant was collected, and Triton X-100 was added to 1%. As input, 5% of the solution was put aside. Primary antibodies: anti-H3K4me1 (abcam, ab8859, 1 µg) anti-H3K4me3 (abcam ab8580, 0.5 μg), anti-H3K9me3 (abcam ab8898, 3.5 μg), anti-H3K27ac (Cell Signaling #8173, 2.5 μg), anti-H3K27me3 (gift from Jenuwein lab, 2.5 μl serum), anti-H3K36me3 (abcam ab9050, 3.5 μg), anti-H2A.xf1^39^ (gift from Shechter lab 2.5 μl serum, anti-H3K79me3 (C15410068, Diagenode, 1 µg), anti-H2AK119ub (Cell Signaling 8240, 2.5 μg). PBS-washed magnetic beads conjugated with secondary antibody (Invitrogen 11204D, 25 μL per 50 embryos) and the fragmented chromatin solution were combined and incubated overnight at 4°C on a rotating wheel. Beads were then washed 6 times with RIPA buffer (50 mM HEPES-KOH pH 7.5, 500 mM LiCl, 1 mM EDTA, 1% Igepal CA-630, 0.7% Na-deoxycholate, complete protease inhibitors (Roche)) and twice with TEN buffer (10mM Tris pH 8.0, 1mM EDTA, 150mM NaCl, complete protease inhibitors (Roche)). For crosslink reversal, the beads were resuspended in stop buffer (40 mM Tris pH 8.0, 10 mM EDTA, 1% SDS). The samples were supplemented with Proteinase K (0.3 μg/μl), NaCl (250 mM) and incubated at 65°C overnight. RNase A (DNase-free) was added to a final concentration of 200 μg/μl and DNA was extracted using phenol/chloroform before 150 μg/μl glycogen was added to recover DNA by ethanol precipitation. The pellet was resuspended in H_2_O.

### ChIP-seq library preparation

The ChIP reaction (above) was subjected for ChIP-seq library preparation with the TruSeq DNA kit (Illumina, FC-121-2001).

### Digital Reprogramming

#### Data pre-processing

For ChIP-seq data reads were first trimmed of their adapters using cutadapt v1.0 and mapped to *Xenopus laevis* genome v9.1 using bwa v0.7.17. Duplicate reads were filtered using samtools v0.1.8 and replicates merged. Peaks were called using the MACS callpeak function vs control Igg levels using the ‘broad’ option where appropriate.

#### Data normalization and augmentation

ChIP-seq data was processed into bigwig format and subsequently into arrays for ease of use in downstream analysis and model inference. For each gene, the histone modification level could be summarised as a (*1 × 600*) length vector by calculating the levels vs Igg in *600* bins running from 5kbp up-stream of the TSS to 5kbp down-stream using the DeepTools computeMatrix function. Additional features, such as DNA methylation levels and %CG content, could similarly be represented as (*1 × 600*) vectors over the same *600* bins. Vectors for all histone modifications and other features in the training set could be concatenated to represented as an (*M × 600*) matrix. Contextually, this can be thought of similarly to a 1D image with *600* pixels and *M* colour channels.

Data was split into training, validation, and test sets, and histone modification levels for donor tissues and target tissues concatenated into arrays: 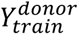 and 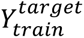 which denote (*N*_*train*_ × *m* × 600) dimensional arrays for the donor and target tissue types respectively, and *M* denotes the number of histone modifications measured. Additionally, 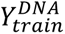 denotes an (*N*_*train*_ × *m* × 600) dimensional array holding the *m* other features e.g., DNA methylation and %CG content. Similarly, all data for test and validation sets could be stored as an arrays 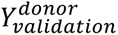, 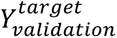, 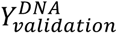, 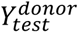, 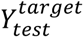, 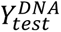. Histone modification levels and other feature levels were row-normalised and scaled to zero-mean unit-variance based upon in training dataset.

Expression levels in donor and target cells could each be represented as a scalar. Thus, for models that use both donor and target tissue expression, this could be represented as *X_train_* = (*N_train_ x 2*) matrix, with corresponding matrices, *X_test_* and *X_validation_* for the test and validation sets. For cases where either the donor or target genes were used for prediction the output is an *X_train_* = (*N_train_ x 1*) column vector, with similar column vectors for the test and validation sets.

Finally, as an output, genes were assigned to one of five gene classes: On-memory (ON), Off-memory (OFF), Reprogrammed-up (RU), Reprogrammed-down (RD), and control regions not belonging to any of the previous four classes (Other). For the control regions, regions from up and downstream of the TSS of genes not belonging to any of the other four classes (On/Off/Up/Down) were randomly sampled. As these classes were mutually exclusive, the output classification for a given gene was indicated via one-hot encoding resulting in a (*1×5*) length vector. Specifically, an On-memory gene was encoded as the output vector [1,0,0,0,0], an Off-memory is [0,1,0,0,0], RU [0,0,1,0,0], RD [0,0,0,1,0], and the control class as [0,0,0,0,1]. The output for all training data is therefore represented as a *Z_train_* = (*N_train_ x 5*) matrix with similar matrices for test and validation sets. The ultimate aim is to predict *Z* given *X*, *Y*.

#### Dimensionality reduction

In order to visualise the data and gauge if histone modification and expression could be used to separate out different classes of genes, we projected processed matrices into lower dimensionality embeddings (PCA, scikit-learn) and via and UMAP using Python.

#### Data augmentation

Due to the limited size of some classes, we opted to augment the dataset to increase the training size and ensure more balanced class representation. Data augmentation was achieved by generating an additional artificial representation of a given gene selected at random by randomly uniformly perturbing the start and end location of the feature from between 1bp and 50bp upstream or downstream. For each randomly perturbed set of chromatin modification features, the corresponding expression scalar was duplicated.

#### Training, test, and validation sets

To ensure no accidental contamination from the training to the test or validation sets e.g., from the data augmentation step, training, test, and validation sets were selected from different chromosomes, with training set corresponding to Chr1-7, test sets to Chr8-14, and validation sets to 15-18.

#### Neural network architectures

Neural network models were coded in keras and consisted of three modules that could be used together or independently, and were loosely inspired by architectures of Angermueller et al.^40^. The first module (<u>Histone Module</u>) integrates chromatin modification and DNA feature data and consisted of three parallel 1D convolution layers with 64 feature maps used to integrate the chromatin data using kernel sizes of 16, 32, and 100 respectively to capture different sized features. Both L1 and L2 regularisation (0.1) and bias were applied to layer weights. Max pooling layers with filter size 4 were applied after convolutional layers. Finally, a dropout layer (0.4) was applied after each max pooling layer. One fully connected (dense) layer with 10 nodes was used to integrate expression data (<u>Expression Module</u>). Histone modules and expression modules were integrated using a fully connected (dense) layer containing 50 and 20 nodes respectively using ReLU activation functions (<u>Integration Module</u>). As we were interested in classifying 1-of-5 categories, the final layer of the integration module consists of 5 nodes with softmax activation, allowing for.

Using these modules, different models were constructed to infer memory gene status as outlined below:

1. **Full (CNN) Model.** The full model combined all three modules to integrate chromatin modification and gene expression levels together. Here we were interested in learning memory status based on both the donor and target tissue histone/gene expression levels i.e., using full knowledge of the state of the gene in both the donor and target tissue. Crucially, this does not make use of any information regarding the state of the reprogrammed state, which is what we aim to infer. Here histone modification arrays for donor and targets were concatenated, 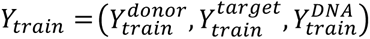, with gene expression concatenated 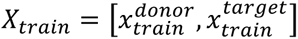. Here we aim to infer *Z*_*train*_|*X*_*train*_, *Y*_*train*_.
2. **Histone only (CMM CNN) Model.** Here we use the histone along with the integration module to infer memory status *Z*_*train*_| *Y*_*train*_. Once again we aim to make full use of the donor and target histone levels, but make no use of the donor and target expression levels.
3. _3._ **A partial (CMM + target CNN) Model.** Here we aim to infer memory status using the histone modification levels of the donor tissues alongside the expression levels of the target tissue type i.e., we are interested in using the histone modification levels in the donor tissues along with knowledge about how the genes are expressed in the target tissue. Thus we have 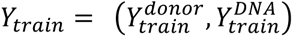, 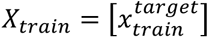, and aim to infer *Z*_*train*_|*X*_*train*_, *Y*_*train*_.
4. **Full (RF) Model.** As a control model, we also included a random forest trained on the full histone modification and gene expression in the donor and target tissues.

For inference, hyperparameters were optimized using SGD optimization with length = 0.001, decay =1e-6, momentum =0.9. Optimization was run for parameters with respect to the categorical cross-entropy for 1000 epochs using a batch size of 1000 genes. Weights were applied to classes inversely proportion to the number of genes in a given class.

### Digital Reprogramming - Transfer learning

Two instances of transfer learning were trialed. In the first, the full neural network e.g., trained using the endoderm to ectoderm data, was used to predict memory status in an alternative condition e.g., the somite to endo data, keeping all parameters (including network architecture, layer weights, and offsets) the same. This transfer learning is referred to as naïve TL. In the second example an updated model was generated by tuning the weights of the densely connected (predictive) layers of the integration module only, but keeping the weights associated with the convolution layers the same. Note that it is the intermediate levels of the convolutional neural network that should be extracting salient features of the histone modification. In the second case, where a degree of optimization was required, inference and pre-processing of datasets were the same as previously highlighted.

### Digital Reprogramming – Accuracy

Area under the curve (AUC) and area under the precision-recall (AUPR) curve were evaluated using sklearn. Other metrics e.g., accuracy calculated using keras during optimization. Average AUC/AUPR were calculated by numerically integrating the AUC/PR curves.

As a control, random forests with sklearn were also used to precinct memory status with n estimators = 100. Here we again inferred memory status using either the histone modification levels, gene expression levels, or both. For use within random forests, histone modification levels were concatenated (along with gene expression levels) e.g., instead of an (*M x B*) matrix for histone modification and (*1 x 2*) for gene expression levels we have a (*1 x MB+2)* length vector for each gene. Output data was again one-hot encoded.

### Digital Reprogramming - Activations

In order to identify putative features that are most likely contributing to memory status, we evaluated layer activations using DeepExplain^43^. Specifically, the Grad * Input was used to calculate activations to ON-memory/OFF-memory/RU/RD classes for a specific set of groups such as true positive genes. We compared these activations to bootstrapped activations to randomly selected outputs.

### p300/CBP inhibitor treatments of donor embryos

IVF-derived gastrulae (NF stage 12) were treated with 40 µM SGC-CBP30 (Sigma, SML1133), 30 µM A-485 (Tocris, #6387) or DMSO in 0.1x MMR at 23°C until they reached neurula stage (NF stage 18)^78^. The embryos were collected at synchronized stages to eliminate influences from different cell numbers or developmental stages. The embryos were either used as donors for nuclear transfer as described above or frozen on dry ice for further experiments.

### Western Blot

Whole embryos were collected at NF stage 18 and lysed in 10 µl E1 buffer (50 mM HEPES-KOH pH 7.5, 140 mM NaCl, 0.1 mM EDTA pH 8.0, 10% glycerol, 0.5% Igepal CA-630, 0.25% Triton X-100, 1% beta-mercaptoethanol) per embryo and then centrifuged at 3500 rpm at 4°C for 2 min to separate the nuclear and cytosolic fraction. For chromatin extraction, the nuclear fraction (pellet) was solubilized in 10 µl E1 buffer per embryo by vigorously agitating and vortexing. Laemmli buffer (BIO-RAD, #1610747) containing 10% beta-mercaptoethanol was added, and the samples were denatured at 95°C for 10 min and centrifuged at maximum speed for 10 min. The chromatin extract was separated on a 4-20% gradient gel (BIO-RAD, Mini-PROTEAN TGX #4561096) and transferred onto a PVDF membrane (Thermo Scientific, 88518) for 1.5 h at 30 V. The membrane was blocked using 5% bovine serum albumin (BSA) and incubated overnight using 1:1000 dilution of primary antibodies against histone modifications (anti-H3K27ac #8173, anti-H3K18ac #9675 from Cell Signaling Technologies; anti-Histone H3 acetyl K9+K14+K18+K23+K27 ab47914 abcam) antibody or anti-histone H4 as a loading control (ab31830, abcam). Protein detection was performed using IRDye-coupled secondary antibodies (IRDye 800CW #926-68022, IRDye 680LT #926-32213, Licor) and imaged on a Licor Odyssey LT machine.

### Histone mass spectrometry and data analysis

50 whole embryos at NF stage 18, pharmacologically treated as described above, were collected in a 2 ml Eppendorf tube, washed 3 times with embryo extraction buffer (10 mM HEPES-KOH pH 7.7, 100 mM KCl, 50 mM sucrose, 1 mM MgCl_2_, 0.1 mM CaCl_2_) by gently inverting the tube, centrifuged at 700 g for 1 min and frozen upon liquid removal at −80°C. To separate the nuclear fraction, the embryos were centrifuged for 10 min at 17000 rpm at 4°C. The nuclei (liquid phase) were carefully removed using a P200 pipette tip, while avoiding the lipids (white ring on top of the liquid phase), and transferred to a fresh Eppendorf tube. The nuclear fraction was washed twice with 500 µl SuNASP (250 mM sucrose, 75 mM NaCl, 0.5 mM spermidine, 0.15 mM spermine) and centrifuged for 5 min at 3500 g at 4°C, without disturbing the nuclear pellet. To extract the histones, the nuclei were solubilized in 600 µl RIPA buffer (50 mM Tris-HCl pH 7.4, 1% Igepal CA-630, 0.25% sodium deoxycholate, 150 mM NaCl, 1 mM EDTA, 0.1% SDS, 0.5 mM DTT, 5 mM sodium butyrate, 1x protease inhibitor) and centrifuged at 14,000 g for 10 min at 4°C. The supernatant was removed, taking care to remove excess lipids or debris. The samples were incubated on ice in 100 µl RIPA, centrifuged as described above and the supernatant was removed. The pellet containing histone extract was solubilized in 50 µl E1 buffer as described for Western Blot, mixed with Laemmli buffer, and separated on a 4-20% gradient polyacrylamide gel. Coomassie-stained bands of H3 and H4 were excised from the gels and stored in Milli-Q H_2_O in 2 ml Eppendorf tubes at 4°C until mass-spectrometry analysis.

Gel pieces containing histones were washed with 100 mM ammonium bicarbonate, dehydrated with acetonitrile, chemically propionylated with propionic anhydride, and digested overnight with trypsin. Tryptic peptides were extracted sequentially with 70% acetonitrile/0.25% TFA and acetonitrile, filtered using C8-StageTips, vacuum concentrated, and reconstituted in 15µl of 0.1% FA.

For LC-MS/MS purposes, desalted peptides were injected in an Ultimate 3000 RSLCnano system (Thermo) and separated in a 25-cm analytical column (75µm ID, 1.6µm C18, IonOpticks) with a 50-min gradient from 2 to 37% acetonitrile in 0.1% formic acid. The effluent from the HPLC was directly electrosprayed into a Qexactive HF (Thermo) operated in data-dependent mode to automatically switch between full scan MS and MS/MS acquisition with the following parameters: survey full scan MS spectra (from m/z 375–1600) were acquired with resolution R=60,000 at m/z 400 (AGC target of 3×106). The 10 most intense peptide ions with charge states between 2 and 5 were sequentially isolated to a target value of 1×105, and fragmented at 27% normalized collision energy. Typical mass spectrometric conditions were: spray voltage, 1.5 kV; no sheath and auxiliary gas flow; heated capillary temperature, 250°C; ion selection threshold, 33.000 counts.

Data analysis was performed with Skyline (v21.2) by using doubly and triply charged peptide masses for extracted ion chromatograms. Automatic selection of peaks was manually curated based on the relative retention times and fragmentation spectra with results from Proteome Discoverer 1.4. Integrated peak values were exported for further calculations. The relative abundance of an observed modified peptide was calculated as the percentage of the overall peptide.

### ChIP-qPCR

ChIP reactions (described above) and input samples were diluted 1:20 and 4 μL were used for subsequent qPCR analysis using primer pairs described in Table 1 at 0.5 µM with iTaq Universal SYBR Green Supermix (BIO-RAD), in a two-step PCR cycle: 94°C for 15 s and 60°C for 60 s. Reactions were performed in a total volume of 20 μl.

**Table 1:**
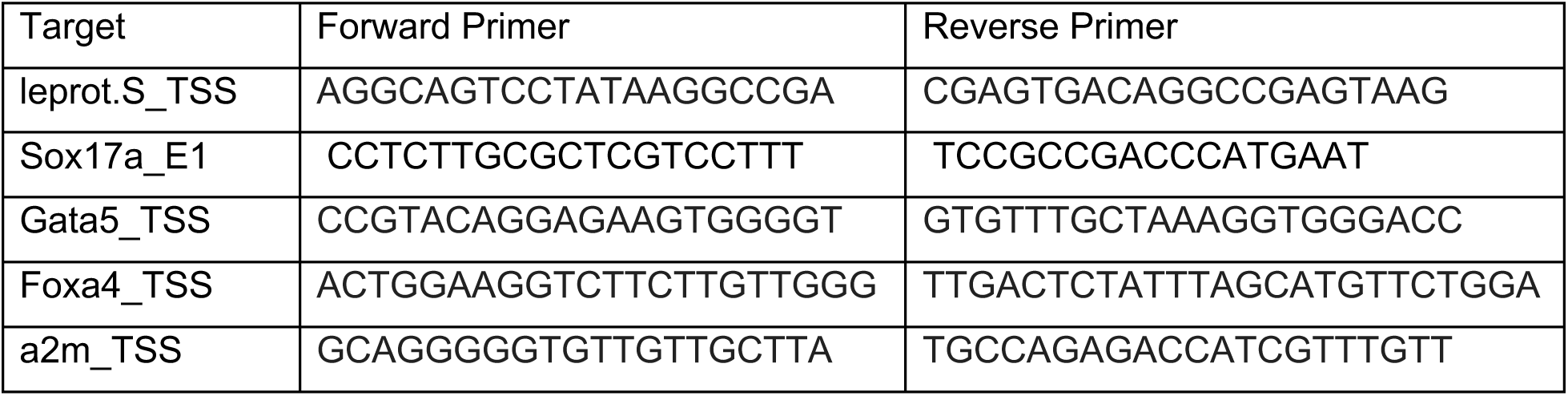
Primers used for ChIP-qPCR assay.

### CUT&RUN

CUT&RUN was performed as described previously^79,80^, with some adjustments. Endoderm was dissected from 25 embryos (NF stage 18) per sample. For nuclear isolation, the endoderm tissues were incubated in Newport 2.0 buffer^81^ to facilitate tissue dissociation. The nuclei were extracted by resuspending in 1 ml ice-cold nuclear extraction buffer (20mM HEPES-KOH, pH 7.9, 10mM KCl, 500 μM spermidine, 0.1% Triton X-100, 20% glycerol), and subsequently resuspended in 600 µl nuclear extraction buffer. 150 µl concanavalin A beads (Epicypher, 21-1401) were resuspended in 850 µl binding buffer (20 mM HEPES-KOH pH 7.9, 10 mM KCl, 1mM CaCl_2_, 1mM MnCl_2_) per sample and activated by washing twice with 1 ml binding buffer. The nuclei were added to 300 µl bead suspension under gentle vortexing and incubated for 10 min at RT while rotating to allow binding of the nuclei to the beads. The supernatant was discarded and the nuclei were incubated in 1 ml blocking buffer (20 mM HEPES-KOH pH 7.5, 150 mM NaCl, 0.5 mM spermidine, 0.1% BSA, 2 mM EDTA,1x protease inhibitor) for 5 min at RT, then incubated in 1:100 primary antibody solution overnight at 4°C and washed twice using 1 ml wash buffer (20 mM HEPES-KOH pH 7.5, 150 mM NaCl, 0.5 mM spermidine, 0.1% BSA, 1x protease inhibitor). To facilitate pAG-MNase (Epicypher, 15-1116) binding to the primary antibody (anti-H3K27ac #8137, anti-IgG #2729; both from Cell Signaling Technologies), samples were incubated under rotation at 4°C for 1 h, washed twice using wash buffer, and resuspended in 150 µl wash buffer. To activate the MNase digestion reaction, 3 µl 100 mM CaCl_2_ was added to each sample and incubated at 0°C for 30 min, then quenched using 2x STOP buffer (200 mM NaCl, 20 mM EDTA, 4 mM EGTA, 50 μg/mL RNase A, 40 μg/mL glycogen), containing 1 ng exogenous spike-in DNA (*E. coli,* Epicypher 18-1401). To release chromatin fragments, the samples were incubated at 37°C for 20 min, following centrifugation at 16.000 g for 5 min at 4°C. The supernatant containing soluble chromatin fragments was mixed with SDS and proteinase K (Sigma-Aldrich) at 70°C for 10 min and subjected to on-column DNA purification (QIAGEN, MiniElute PCR Purification Kit).

### CUT&RUN sequencing library preparation

CUT&RUN sequencing libraries were prepared using NEB Ultra II DNA library prep kit (NEB, #E7645), following the manufacturer’s instructions, using 12 PCR amplification cycles with CUT&RUN-specific cycling parameters (45 s 98°C for polymerase activation, 14 cycles of 15 s 98°C DNA melting and 10 s 60°C primer annealing and short extension, 1 min 72°C final extension), without size selection. The libraries were sequenced on a NovaSeq X+ sequencing platform.

### Experimental design

In all experiments analyzing gene expression, one sample corresponds to tissue from one embryo. In the experiments addressing the effect of p300/CBP expression using CBP30 on ON-memory in NT-embryos, 3 independent experiments (referred to as biological replicates) were performed. In each experiment, 2 endoderm-donor samples were collected per condition: donor(DMSO) and donor(CBP30) and 4 IVF-ectoderm samples. In experiment #1, 4 NT(DMSO)-ectoderm samples and 3 NT(CBP)-ectoderm samples were collected. In experiment #2, 3 NT(DMSO)-ectoderm samples and 5 NT(CBP)-ectoderm samples were collected. In experiment #3, 5 NT(DMSO)-ectoderm samples and 3 NT(CBP)-ectoderm samples were collected.

For the A-485 analysis, 3 independent experiments were performed, designed as described for CBP30. As a control, the inactive compound A-486 was used. In each experiment, 2 endoderm-donor samples were collected per condition: donor(A-486) and donor(A-485) and 4 IVF-ectoderm samples. In experiment #1, 5 NT(A-486)-ectoderm samples and 3 NT(A-485)-ectoderm samples were collected. In experiment #2, 5 NT(A-486)-ectoderm samples and 5 NT(A-485)-ectoderm samples were collected. In experiment #3, 3 NT(A-486)-ectoderm samples and 3 NT(A-485)-ectoderm samples were collected.

For histone mass-spectrometry analysis, three independent experiments were performed (referred to as biological replicates), in which DMSO, CBP30, A-485, A-486 and untreated embryos were collected in parallel for each biological replicate.

For CUT&RUN, four independent experiments were performed, in which endoderm tissue was collected from DMSO, CBP30 and A-485 treated embryos, in parallel for each biological replicate. Endoderm tissue was collected from 25 embryos per condition, per sample, and pooled for nuclear extraction.

### RNA-seq Data Processing and Differential Expression Analysis

Paired sequencing reads were processed using Kallisto (v0.48) for pseudoalignment and quantification of transcript abundance. Transcript and annotation files were downloaded from Xenbase (v10.1) for *Xenopus laevis* (transcripts and annotation). After quantification, transcript-level abundances were imported using ‘tximport’ (v1.32.0) and converted to a ‘SummarizedExperiment’ (v1.34.0) object in R (v4.3.1). Datasets of two independent batches were merged and subjected to differential expression analysis performed with DESeq2 (v1.40.2).

### Filtering strategy for memory class and reprogrammed genes

For RNA-seq experiments addressing the effects of p300/CBP inhibition on ON-memory, log2 fold changes (log2FC) and adjusted p-values (p-adj) were calculated using DEseq2^82^. The gene lists were then filtered as follows (note that 3FC corresponds to log2FC ≈ 1.5).

<u>Differentially expressed genes (DEG) Donor/IVF:</u> p-adj (Donor/IVF) < 0.05 DEG Donor/IVF and NT/IVF: p-adj (Donor/IVF) < 0.05 & p-adj (NT/IVF) < 0.05

<u>ON-memory genes:</u> p-adj (Donor/IVF) < 0.05, log2FC (Donor/IVF) > 0, p-adj (NT/IVF) < 0.05, log2FC(NT/IVF) > 0, TPM(Donor) >1.

<u>ON-memory (3FC) genes:</u> p-adj (Donor/IVF) < 0.05, log2FC (Donor/IVF) > 0, p-adj (NT/IVF) < 0.05, log2FC(NT/IVF) > 1.5, TPM(Donor) >1.

<u>OFF-memory genes:</u> p-adj (Donor/IVF) < 0.05, log2FC (Donor/IVF) < 0, p-adj (NT/IVF) < 0.05, log2FC(NT/IVF) < 0

<u>OFF-memory (3FC) genes:</u> p-adj (Donor/IVF) < 0.05, log2FC (Donor/IVF) < 0, p-adj (NT/IVF) < 0.05, log2FC(NT/IVF) < 1.5

<u>Reprogrammed-Down</u>: p-adj (Donor/IVF) < 0.05, log2FC (Donor/IVF) >0, p-adj (Donor/NT) < 0.05, log2FC (Donor/NT) > 0.05, TPM (Donor) >1. Transcripts with p-adj (NT/IVF) < 0.05 were excluded.

<u>Reprogrammed-Up:</u> p-adj (Donor/IVF) < 0.05, log2FC (Donor/IVF) < 0, p-adj (Donor/NT) < 0.05, log2FC (Donor/NT) < 0.05, p-adj (NT/IVF) > 0.05.

In the experiments comparing the effect of p300/CBP inhibitor treatment vs. control, filtering the ON-memory genes was performed separately in each condition.

<u>Differentially expressed genes (DEG) Donor(DMSO)/IVF:</u> p-adj (Donor(DMSO)/IVF) < 0.05

<u>DEG Donor(DMSO)/IVF and NT(DMSO)/IVF:</u> p-adj (Donor(DMSO)/IVF) < 0.05 & p-adj (NT(DMSO)/IVF) < 0.05

<u>ON-memory genes (DMSO):</u> p-adj (Donor(DMSO)/IVF) < 0.05, log2FC (Donor(DMSO)/IVF) > 0, p-adj (NT(DMSO)/IVF) < 0.05, log2FC(NT(DMSO)/IVF) > 0, TPM(Donor(DMSO)) >1.

<u>ON-memory (3FC) genes (DMSO):</u> p-adj (Donor(DMSO)/IVF) < 0.05, log2FC (Donor(DMSO)/IVF) > 0, p-adj (NT(DMSO)/IVF) < 0.05, log2FC (NT(DMSO)/IVF) > 1.5, TPM(Donor(DMSO)) >1.

<u>OFF-memory genes (DMSO):</u> p-adj (Donor(DMSO)/IVF) < 0.05, log2FC (Donor(DMSO)/IVF) < 0, p-adj (NT(DMSO)/IVF) < 0.05, log2FC(NT(DMSO)/IVF) < 0

<u>OFF-memory (3FC) genes (DMSO):</u> p-adj (Donor(DMSO)/IVF) < 0.05, log2FC (Donor(DMSO)/IVF) < 0, p-adj (NT(DMSO)/IVF) < 0.05, log2FC(NT(DMSO)/IVF) < 1.5

<u>Reprogrammed-Down (DMSO)</u>: p-adj (Donor(DMSO)/IVF) < 0.05, log2FC (Donor(DMSO)/IVF) >0, p-adj (Donor(DMSO)/NT(DMSO)) < 0.05, log2FC (Donor(DMSO)/NT(DMSO)) > 0.05, TPM (Donor(DMSO)) >1. Transcripts with p-adj (NT(DMSO)/IVF) < 0.05 were excluded.

<u>Reprogrammed-Up (DMSO):</u> p-adj (Donor(DMSO)/IVF) < 0.05, log2FC (Donor(DMSO)/IVF) < 0, p-adj (Donor(DMSO)/NT(DMSO)) < 0.05, log2FC (Donor(DMSO)/NT(DMSO)) < 0.05, p-adj (NT(DMSO)/IVF) > 0.05.

The same filtering strategy was applied for samples generated upon CBP30 and A-485 treatment.

### Plots for gene expression

Heatmaps – Expression values for each condition were normalized to the mean expression of IVF controls. Log2-transformed fold changes were calculated for each condition relative to IVF mean expression. These values were plotted using heatmap.2 from the R package gplots (v 3.2.0), with clustering by rows and columns and default settings: complete as agglomeration method and Euclidean distance as similarity measure. Z-scores were calculated per row, using the scaling function in heatmap.2 (v3.2.0).

MA-plots – The log2FC between genes expressed in NT and IVFs was plotted against the log2-transformed TPM-expression values (log2 TPM+1) in the endoderm donor cells. MA-plots were created using base R.

Boxplots – The distribution of mean expression levels of the different gene sets is represented as boxplots, whereby the line represents the median, the box edges indicate the lower and upper percentiles (25^th^ and 75^th^ respectively), and the whiskers represent the minimum and maximum values. Boxplots were created using default settings in base R.

p-values comparing differences in gene expression between the different gene sets were calculated using a two-sided unpaired Wilcoxon rank-sum test in R.

Principal component analysis (PCA) plots - Principal Component Analysis was performed on TPM gene expression data. Genes with zero variance were removed. The data was centered and scaled prior to PCA. Finally, PCA was performed using the prcomp(v3.6.2) package and visualized using ggplot2(v. 3.5.1) in R.

### ChIP-seq analysis for enhancer annotation

Raw ChIP-seq data were aligned to the *Xenopus laevis* genome (v10.1) using BWA (v0.7.17-foss-2018b). Low-quality mapped reads (MAPQ < 10) were discarded using samtools (v1.16.1), and PCR duplicates were removed with sambamba (v0.8.2). Coverage track files (.bigwig) were generated using deepTools (v3.5.1), with normalization to Counts Per Million (CPM). Overlaps between peaks and genes were analyzed using the GenomicRanges^83^ package (v1.52.1). Heatmaps were created in R (v4.3.0) using the EnrichedHeatmap^84^ package (v1.30.0).

### Plots for enrichment of individual histone modifications

The coverages were analyzed using the EnrichedHeatmap^84^ package for three distinct genomic contexts: promoters, gene bodies, and putative enhancers. Promoter regions were defined as ±1kb windows centered on the TSS. For the enhancer analysis, we identified putative enhancer regions by p300 binding peaks, excluding any regions intersecting with promoter windows. For assigning putative enhancer regions to target genes, we used our custom-made annotation following Ing-Simmons et al.^55^ and as described below. Coverage values were aggregated for genes with multiple enhancer associations by summing the signals.

The resulting coverages were log2 transformed after adding a pseudo count of 1 to provide the ChIP-seq intensities used for meta-plots and heatmaps.

Meta plots (line plots) – Aggregate line plots, depicting the mean ChIP-Seq intensity of gene sets of interest with bands representing standard error of the mean, were generated using ggplot2.

Heatmaps – Heatmaps were generated using the EnrichedHeatmap package in R. Each row represents the ChIP-Seq intensity of a given histone mark on one gene.

### ChIP-seq analysis for enhancer annotation

For the endoderm H3K27ac ChIP-seq analysis, peaks were called using MACS2^85^ (v2.2.8) with the “--broad” option. Due to low enrichment and minimal peak calls, replicate 2 was excluded from further analysis. Common peaks between replicates 1 and 3 were identified and merged using bedtools (v2.31.0), resulting in a total of 7915 peaks. These peaks were further classified into 4851 promoter peaks (+/-500bp) and 3064 non-promoter peaks (distant from promoters by more than 500bp). To predict active enhancers in *Xenopus laevis* endoderm, previously published p300 ChIP-seq data^54^ from NF stage 20 foregut and hindgut were analyzed. Foregut and hindgut peaks were concatenated (cat), sorted (sort -k1,1 -k2,2n) and merged using bedtools merge, resulting in 61418 peaks on the main chromosomes. Subsequently, we identified 1306 H3K27ac+ and p300+ putative enhancers, supported by detected H3K4me1 signal on these sites. Enhancer-gene pairing was performed as described previously^55^.

### CUT&RUN analysis

Sequencing reads were aligned to the *Xenopus laevis* genome (v10.1) and a spike-in genome (*Escherichia coli*, GCA_001606525) using Bowtie2 (v2.4). The alignment process excluded discordant and mixed reads, with paired-end reads mapped to the reference genome. Aligned reads were converted to BAM format and filtered to retain only properly paired alignments. Normalization scaling factors were computed using deepTools (v3.5.3) based on the read distribution across genomic regions. HOMER (v4.11) was used for peak identification. Broad peaks were called using histone-style parameters and converted to BED format for downstream analysis. Differential peak analysis for H3K27ac signal was performed on the p300 ChIP-seq peak set from Stevens et al.^54^ as a reference peak set using DEseq2^82^ (v1.46.0). Enhancers and promoters were annotated as described above.

Boxplots comparing DEseq2 normalized counts on differentially abundant peaks across gene sets of interest were generated in base R (v4.4.2), whereby the line represents the median, the box edges indicate the lower and upper percentiles (25^th^ and 75^th^ respectively), and the whiskers represent the minimum and maximum values. P-values for pairwise comparisons were calculated using a two-sided Wilcoxon rank-sum test.

MA-plots, balloon plots, stacked bar charts, and scatter plots were generated using ggplot2 (v3.5.1) in R. Heatmaps comparing enrichment or correlations were generated using pheatmap (v1.0.12) in R.

Heatmaps comparing global coverage distribution across conditions were generated in R using the package EnrichedHeatmap^84^, using the function normalizeToMatrix to calculate the coverage over genomic features. The coverage from biological replicates was averaged and normalize to *E. coli* spike-in.

Enrichment testing was performed in R using a two-sided Fisher’s text and p-values were corrected for multiple comparisons using the Benjamini-Hochberg method.

## Data Availability

Raw data from ChIP-sequencing, RNA-sequencing and CUT&RUN are available at GEO under accessions GSE285205, GSE285204 and GSE285202, respectively. Code used for analysis in this paper is available at the GitHub repository https://github.com/cap76/DeepXen. Raw data from histone mass-spectrometry have been deposited to the ProteomeXchange^86^ Consortium via the PRIDE^87^ partner repository with the dataset identifier PXD056817.

## ACKNOWLEDGEMENTS

Work in the E.H. lab was supported by CRC1064 Chromatin Dynamics Project-ID 213249687 and HO 6864/2-1 project grant, both DFG; Recognition Award, Helmholtz Association; Marie Sklodowska-Curie Postdoctoral Fellowship; funding from the Helmholtz Association. Work in the JBG lab was funded by the Molecular Research Council (MR/P000479/1), the Wellcome Trust (101050/Z/13/Z and 092096/Z/10/Z), and Cancer Research UK (C6946/A14492). M.S. received funding from the Joachim Herz Stiftung and “Munich School for Data Science— MUDS”. T.S. lab was funded DFG (CRC1064/Z04). The authors are grateful to the members of the CAM LMU Munich, Dave Simpson and Rue Green for supporting frog work. We thank all present and past lab and institute members, especially Meghana Oak, Antonio Scialdone and Maria-Elena Torres-Padilla, as well as Ralph Rupp and Kikuë Tachibana for reagents, their critical input and support.

## Author contributions

A.J., C.A.P., J.J., S.L.A., H.L., M.S., L.H., T.S., T.Z., I.F., and E.H. performed experiments and analyzed the data. A.J, C.A.P, J.J. and E.H. designed experiments and analyses. J.J. and E.H. obtained funding, A.J, C.A.P and E.H. wrote the paper. A.J., C.A.P., J.J., S.L.A., H.L., M.S., L.H., T.S., T.Z., I.F. and E.H. reviewed and edited the paper.

## SUPPLEMENTARY MATERIALS – TABLES

**Table S1:**
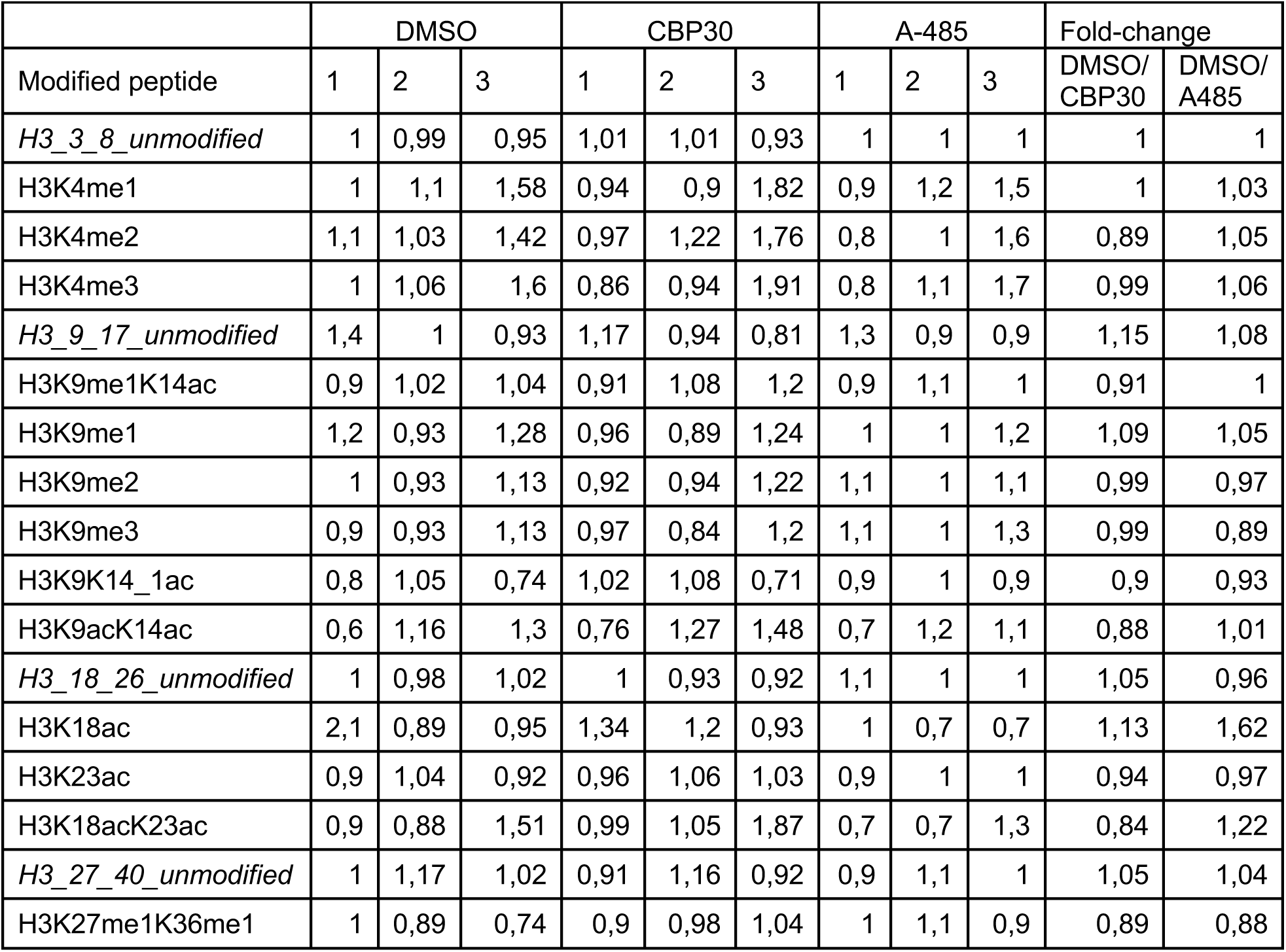
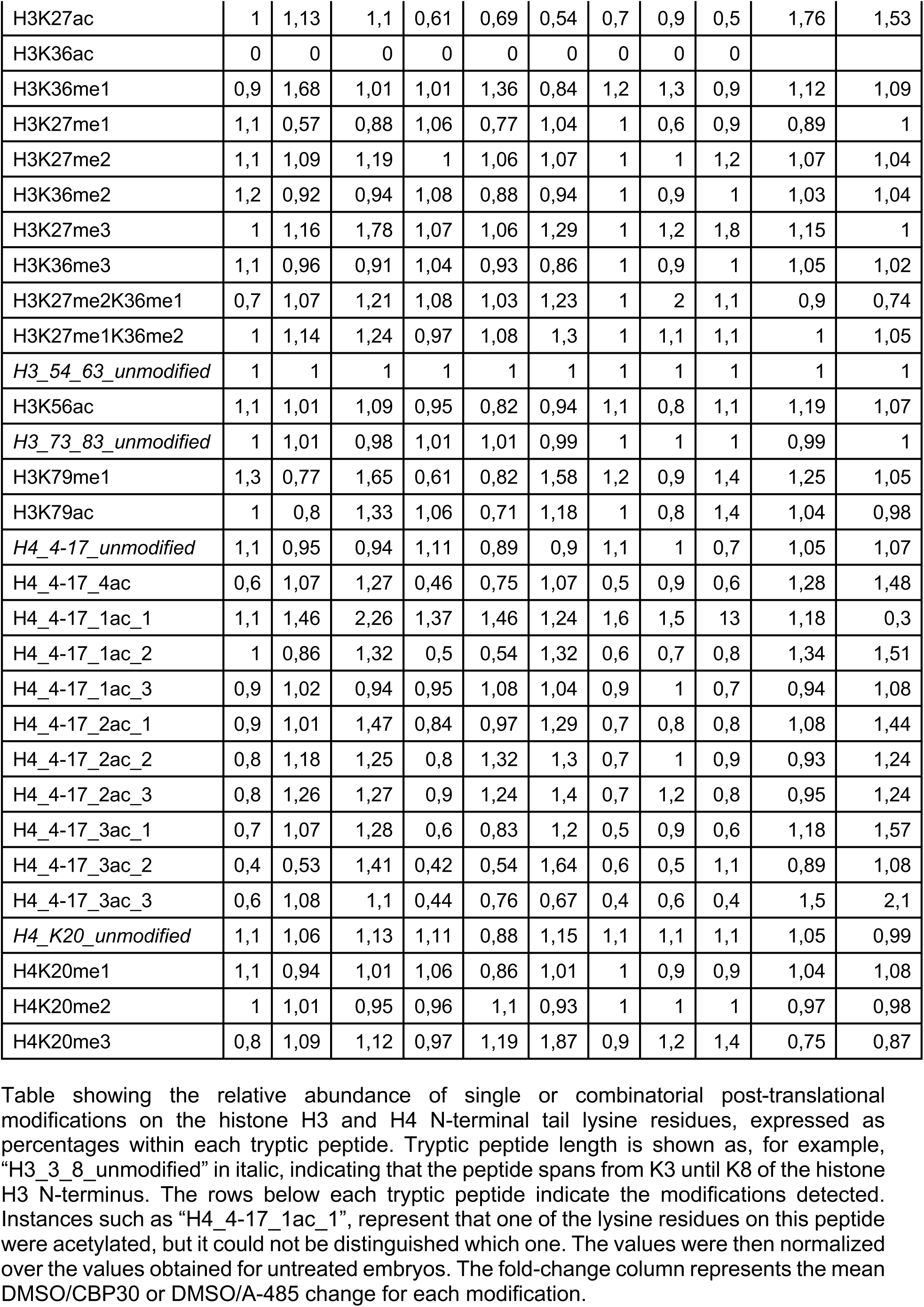
Histone Tail mass-spectrometry data in DMSO, CBP30 and A-485 samples, related to Figure 3.

**Table S2:**
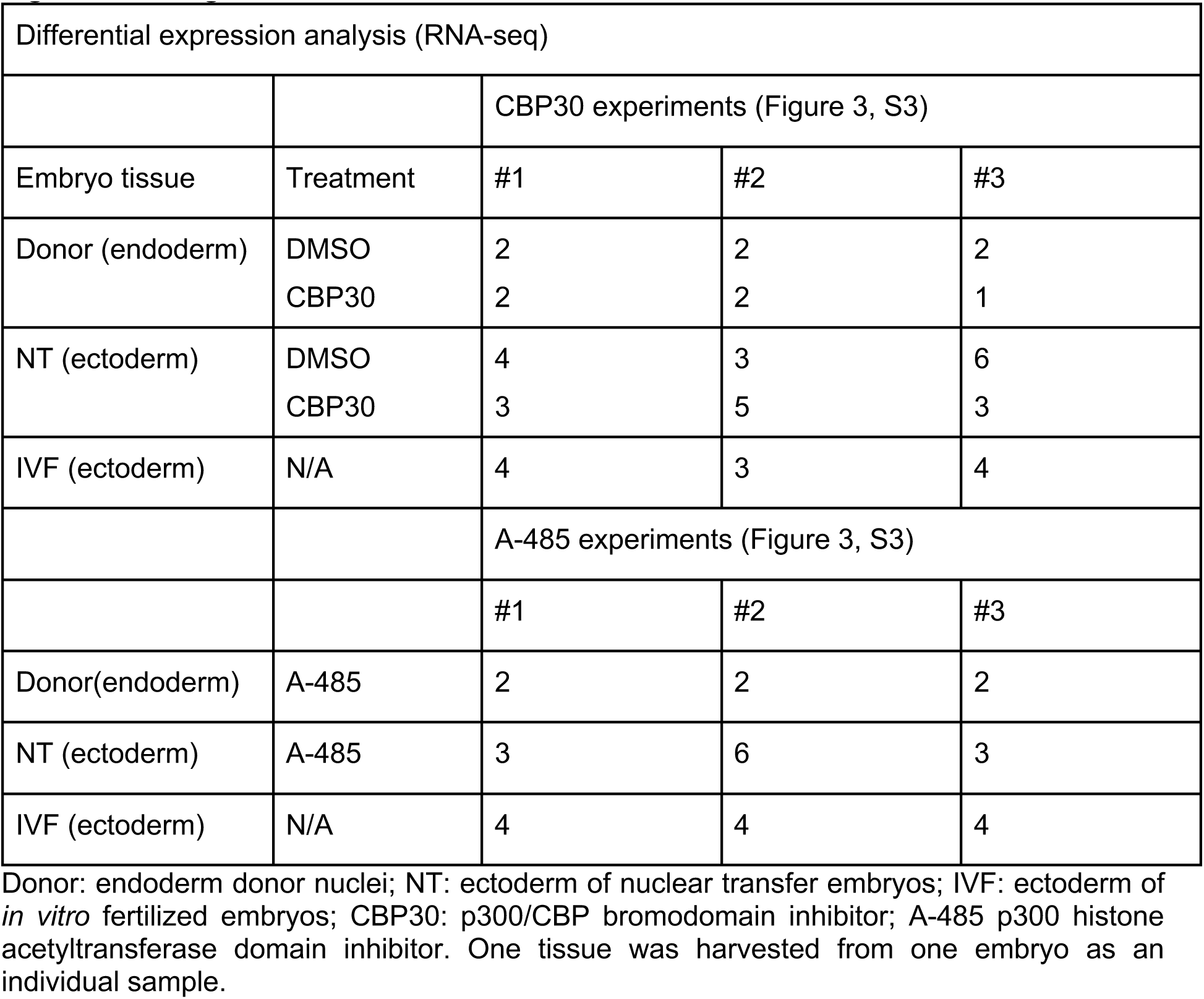
Overview of Biological Replicates in transcriptome experiments related to Figure 3 and Figure S3.

**Table S3:** Overview of gene expression changes in p300/CBP inhibitor treated donor cells (CBP30 - bromodomain inhibitor, or A-485 - catalytic domain inhibitor)

|  | Number of transcripts | % of transcripts | Filter applied |
| --- | --- | --- | --- |
| Total transcripts | 36994 | 100 | TPM >1 in either donor, IVF or NT samples |
| DE donor DMSO vs. donor CBP30 | 3282 | 8,8 | p-adj < 0.05 |
| DE donor DMSO vs. donor A-485 | 5493 | 14,8 | p-adj < 0.05 |

Donor: endoderm donor nuclei; NT: ectoderm of nuclear transfer embryos; IVF: ectoderm of *in vitro* fertilized embryos; DE: differentially expressed; TPM: transcripts per million; p-adj: adjusted p-value.

**Table S4:** Overview of memory and reprogrammed gene sets upon filtering.

|  | DMSO |  | CBP30 |  | A-485 |  |
| --- | --- | --- | --- | --- | --- | --- |
|  | Number of transcripts | % of total transcripts | Number of transcripts | % of total transcripts | Number of transcripts | % of total transcripts |
| Total transcripts | 36994 | 100 | 36994 | 100 | 36994 | 100 |
| DE Donor vs. IVF | 20314 | 54,9 | 19336 | 52,2 | 19379 | 52,4 |
| DE Donor vs. IVF & DE NT | 17662 | 47,7 | 16773 | 45,3 | 16456 | 44,5 |
| ON-memory | 1360 | 3,6 | 1071 | 2,9 | 1253 | 3,4 |
| OFF-memory | 1315 | 3,55 | 1413 | 3,8 | 1525 | 4,1 |
| RD | 5906 | 15,9 | 5186 | 14,0 | 5153 | 13,9 |
| RU | 3754 | 10,1 | 6223 | 16,8 | 5787 | 15,6 |

Donor: endoderm donor nuclei; NT: ectoderm of nuclear transfer embryos; IVF: ectoderm of *in vitro* fertilized embryos; DE: differentially expressed; RD: reprogrammed-down genes; RU: reprogrammed-up genes. The filtering strategy is described in the Materials and Methods section

**Table S5:**
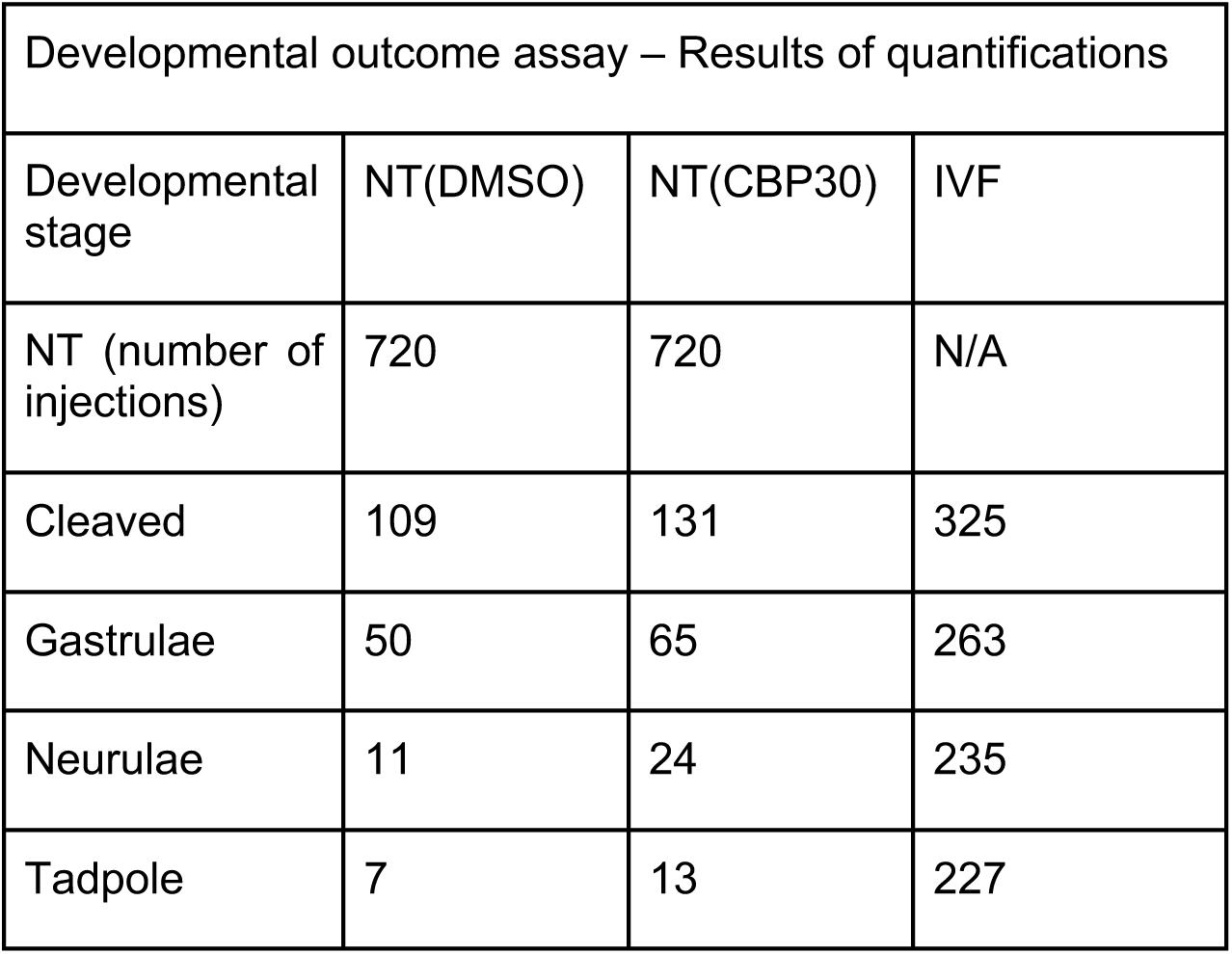
Overview of embryo counts in the developmental outcome assay (Fig. 3, S3)

## Supplementary figure legends

**Figure S1:**
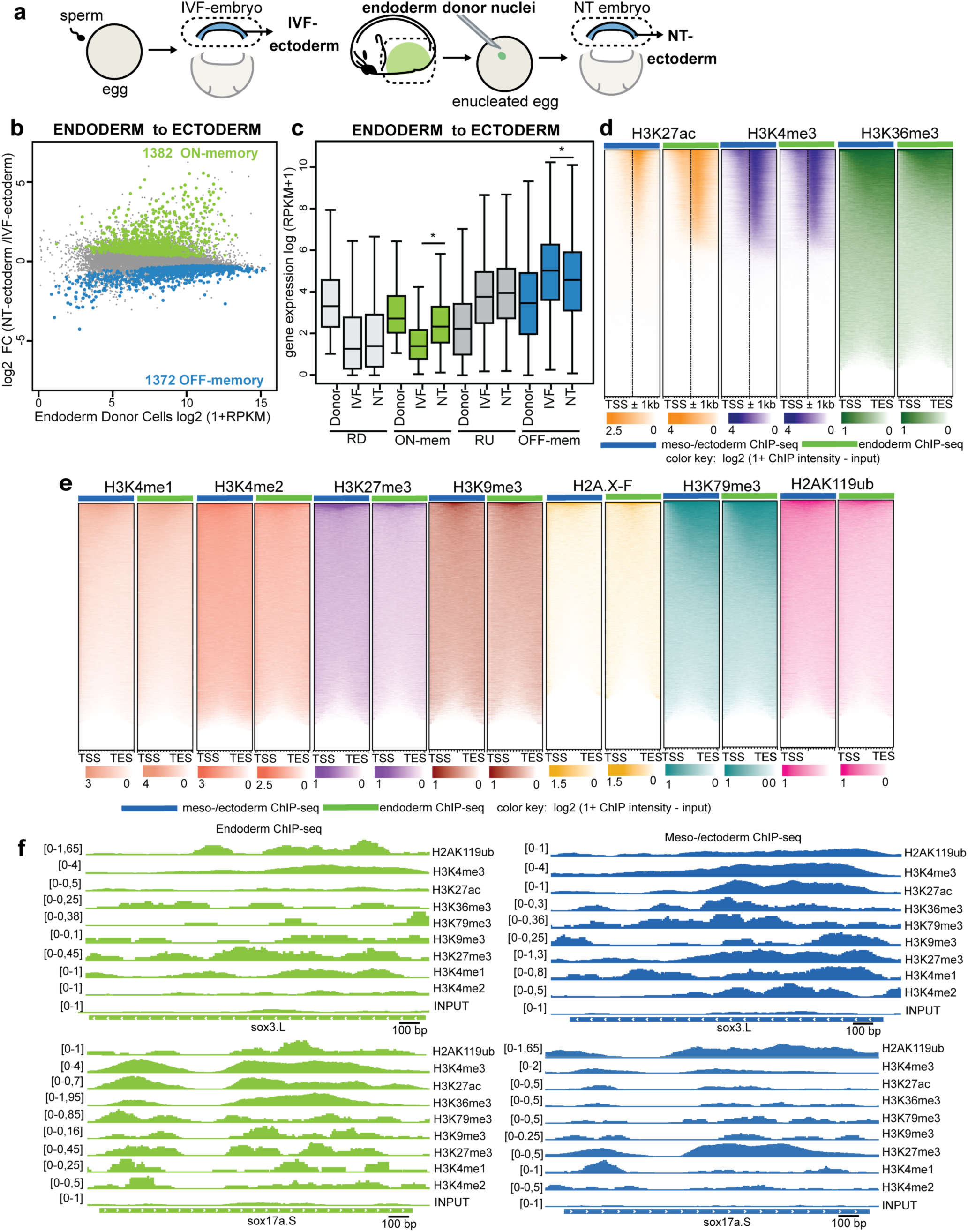
Memory genes resist reprogramming. **(a)** Memory genes were identified in stage 11 ectoderm following NT with endoderm donor nuclei compared to IVF ectoderm. **(b)** MA-plot comparing gene expression between ectoderm samples in NT versus IVFs. The mean log2-fold change gene expression in NT-embryos over IVF is plotted on the y-axis, while the mean log2(RPKM+1) gene expression in endoderm donor nuclei is plotted on the x-axis. NT reprogramming from endoderm to ectoderm revealed 1382 ON-memory genes (green) and 1372 OFF-memory genes (blue). **(c)** Boxplots showing log2(RPKM+1) mean expression levels of Reprogrammed-Down (RD; light gray), ON-memory (ON-mem; green), Reprogrammed-Up (RU; dark gray) and OFF-memory (OFF-mem; blue) genes in endoderm donor samples and ectoderm IVF and NT samples. p-values for pairwise comparisons were calculated using Wilcoxon rank-sum test **(d-e)** Heatmaps representing ChIP-seq signal distribution around the TSS (+/-1 kb) for **(d)** H3K4me3 and H3K27ac and gene body (from TSS to TES) for H3K36me3, as well as **(d)** H3K27me3, H3K9me3, H3K36me3, H3K79me3, H3K4me1 and H2AK119ub. Color intensity represents 1+log2 transformed coverage for each histone mark. Rows represent individual genes on the main chromosomes of *X. laevis*. Green bars indicate ChIP-seq samples from endoderm, while blue bars indicate meso-/ectoderm samples. **(d)** Genome browser snapshots representing histone mark enrichment on candidate genes (*sox3.L* and *sox17a.S*) in endoderm and meso-/ectoderm samples.

**Figure S2:**
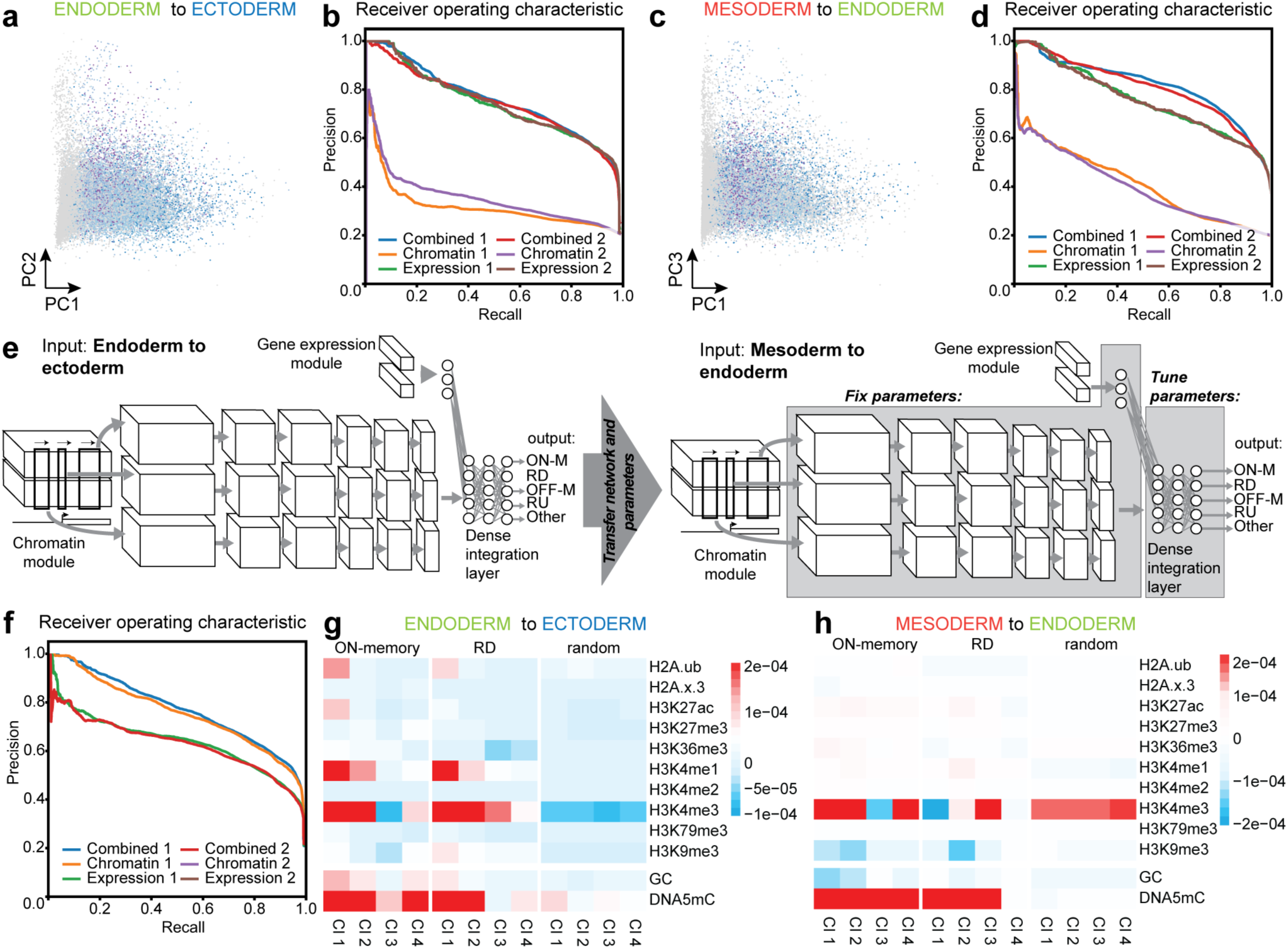
Digital Reprogramming predicts Memory genes in NT reprogramming and identifies novel epigenetic barriers. **(a)** PCA representations of memory class genes projected by histone modification level and gene expression for endoderm to ectoderm reprogramming. **(b)** Receiver operating characteristic (ROC) curve, endoderm to ectoderm reprogramming. **(c)** PCA representations of memory class genes projected by histone modification level and gene expression, mesoderm to endoderm reprogramming. **(d)** ROC curve for mesoderm to endoderm reprogramming. **(d)** Schematic for full transfer learning, including optimization of lower levels of the CNN, used to gauge the effectiveness of learned features. **(d)** Receiver operator characteristic (ROC) curves for transfer learning (d). **(g and h**) Average activations of ON-memory genes for ON-memory, reprogrammed-down and random in endoderm to ectoderm (g) and mesoderm to endoderm reprogramming (h). Color indicates the sum over the cluster average activation over 600-bins of the TSS-flanking region, with higher scores indicating a greater contribution of that histone modification to predicted memory status.

**Figure S3:**
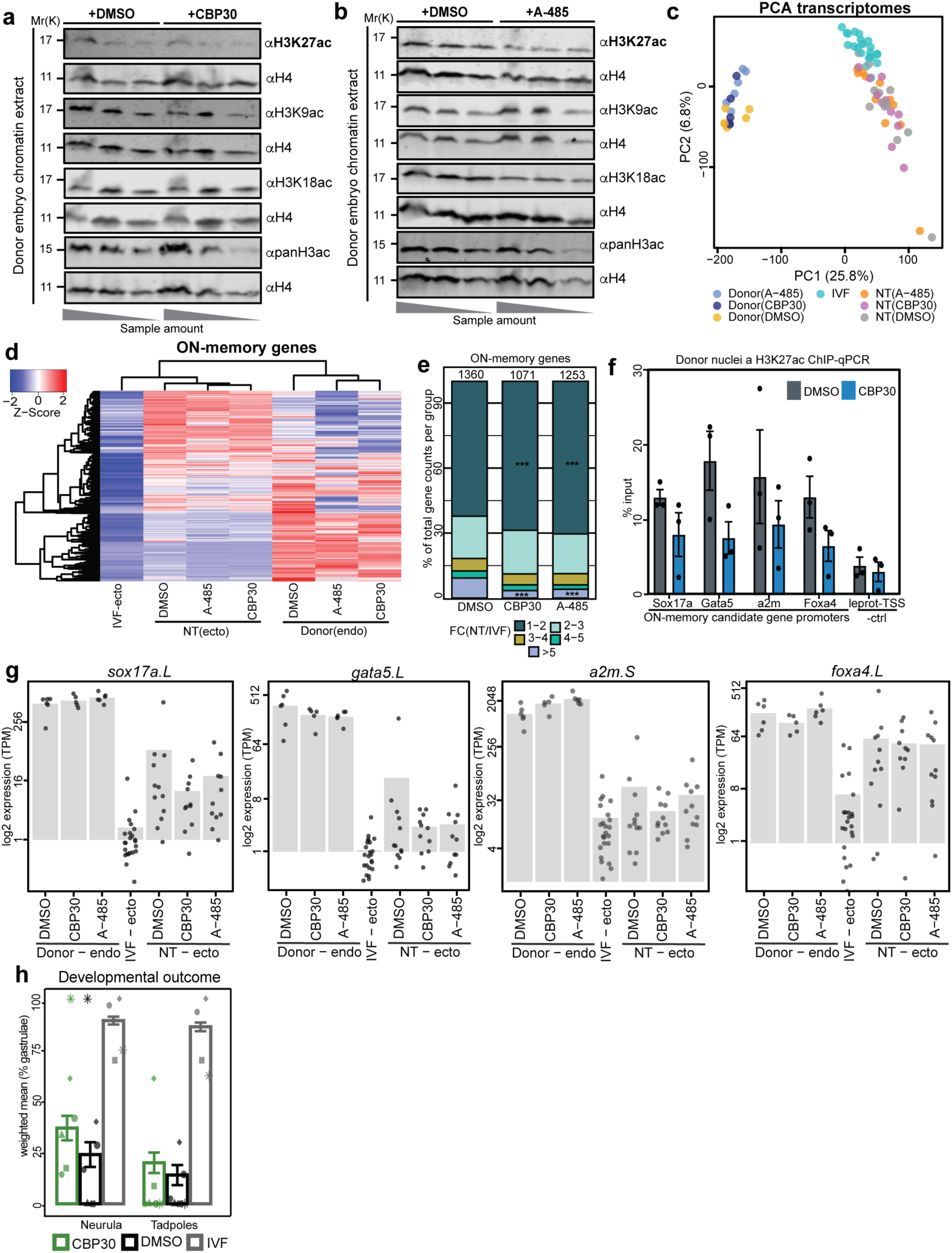
p300/CBP perturbation in donor nuclei reduces histone acetylation and improves ON-memory in NT-embryos. **(a-b)** Western Blot analysis of H3K27ac, H3K9ac, H3K18ac and pan-H3-acetylation (acetyl K9+K14+K18+K23+K27) levels for (a) CBP30 treated donor nuclei vs. DMSO control and (b) A-485 treated donor nuclei vs. DMSO control. Histone H4 was used as a loading control. Each sample was loaded as a dilution series in 2-fold dilution steps, from the highest to the lowest sample concentration. **(c)** PCA comparing the total transcriptome of Donor(DMSO), Donor(CBP30), Donor(A-485), IVF, NT(DMSO), NT(CBP30) and NT(A-485) samples. X-axis represents Principal Component (PC) 1, explaining 25,8% of the variance, y-axis represents PC2 explaining 6,8% of the variance. **(d)** Heatmap representing relative gene expression levels compared to mean IVF expression in endoderm donor samples treated with DMSO, CBP30 or A-485, IVF ectoderm, and NT ectoderm samples derived from donors treated with DMSO, CBP30 or A-485. Rows and columns were clustered using hierarchical clustering and Euclidean distance. The color key represents the z-score on rows. **(d)** Percentage of ON-memory genes defined under DMSO (n=1360), CBP30 (n=1071) or A-485 (n=1253) conditions, stratified based on their fold change in NT/IVF samples. two-sided Fisher’s test, *** p-value < 0.0001. **(f)** ChIP-qPCR targeting regions around the promoter of *sox17a.L*, *gata5.L*, *a2m.S* and *foxa4.L* in DMSO and CBP30-treated samples. y-axis: H3K27ac levels as % input. *leprot.S*-TSS serves as a negative control locus. **(g)** Bar charts representing mean expression levels (log2(TPM)) of candidate ON-memory genes *sox17a.L*, *gata5.L*, *a2m.S* and *foxa4.L*. **(h)** Bar charts representing the weighted mean +/-SEM of obtained NT(DMSO), NT(CBP30) and IVF embryos, expressed as a percentage of gastrulated embryos. Each data point represents an independent experiment, N=6.

**Figure S4:**
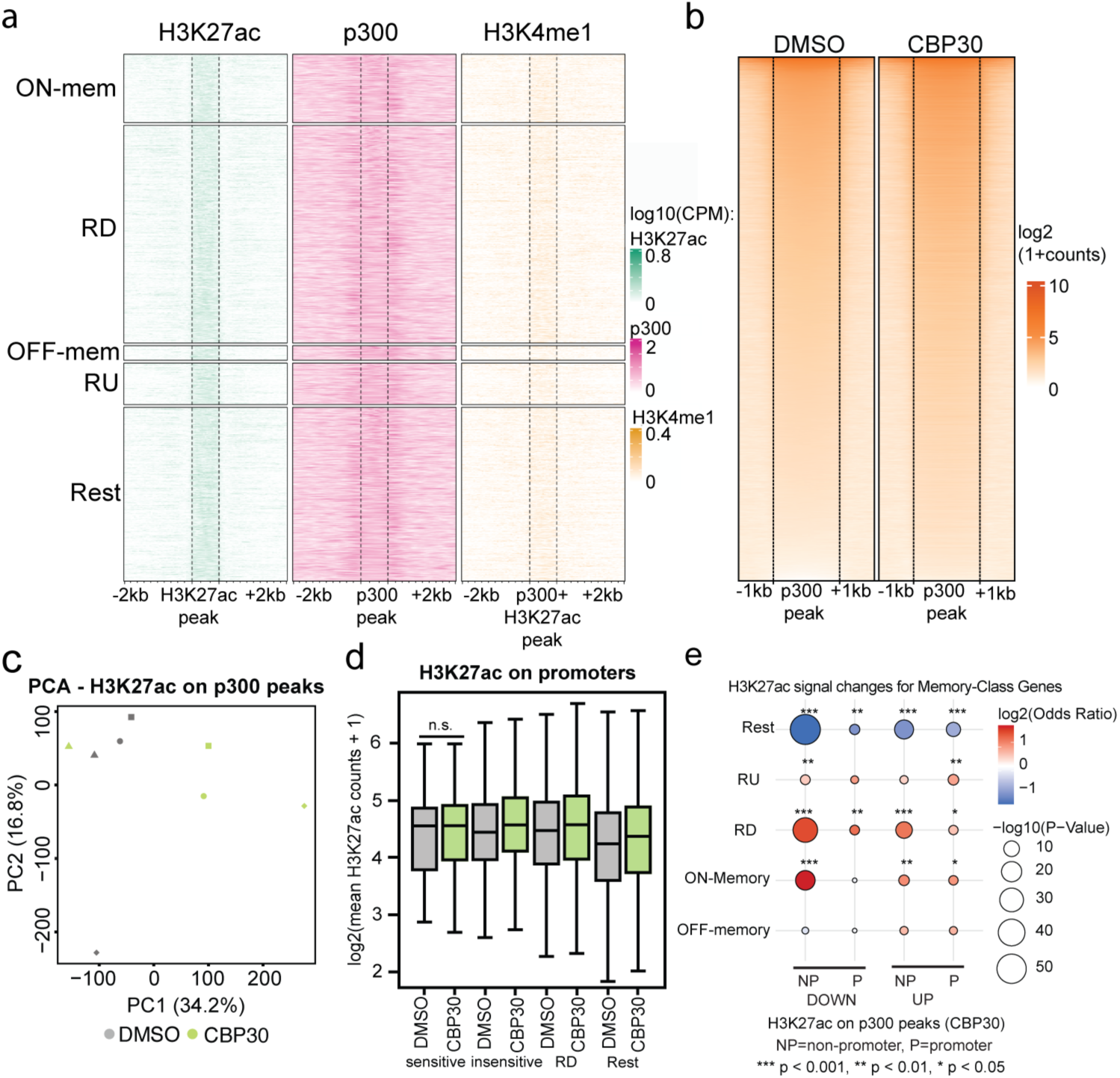
ON-memory genes are enriched for active enhancers, which are targeted by CBP30 and demonstrate reduced H3K27ac levels. **(a)**Heatmap showing H3K27ac, p300 and H3K4me1 signal (log2 counts per million, CPM) on predicted enhancers, clustered for ON-memory, OFF-memory, reprogrammed-down and reprogrammed-up genes. **(b)** Heatmaps showing mean H3K27ac signal (log2 counts+1) across 4 biological replicates detected via CUT&RUN in DMSO and A-485 samples, centered around p300-peaks including promoters and enhancer peaks +/-1 kb. **(c)** Principal component analysis comparing H3K27ac counts on p300-peaks in DMSO (gray) vs. CBP30 (green) samples, *E. coli* spike-in normalized. **(d)** Balloon plot comparing enrichment of memory class and reprogrammed gene sets versus changes in H3K27ac levels on promoters or non-promoter p300-peaks. Balloon sizes represent the -log10 (p-value) and colors represent log2(odds ratio). **(e)** Boxplot comparing H3K27ac levels between donor(DMSO) and donor(CBP30) on promoters proximal to treatment-sensitive ON-memory genes (n= 23 peaks, p-value = n.s.), treatment-insensitive ON-memory genes (n= 741, p-value = 0.001), RD genes (n= 2654, p-value = 8.32·10^-6^), and Rest (n= 8993, p-value = 1.67·10^-20^).

